# A field experiment assessing the roles of drought, herbivory, and local climate on cyanogenesis cline formation and local adaptation in *Trifolium repens*

**DOI:** 10.1101/2025.03.07.642096

**Authors:** Lucas J. Albano, Courtney M. Patterson, Evan J. Kidder, Nicholas J. Kooyers, Marc T. J. Johnson

**Author notes:** **Correspondence Author:** Lucas J. Albano, Current address: Department of Biology, University of Toronto Mississauga, Mississauga, ON, Canada, L5L 1C6.

## Abstract

Projecting how a population will adapt to environmental changes requires a mechanistic understanding of the specific biotic and abiotic factors that impose selection on that population. In white clover (*Trifolium repens*), clines in an antiherbivore defense mechanism, hydrogen cyanide (HCN), form via variation in selection imposed by the environment. However, the specific environmental factors that select for or against chemical defense phenotypes in white clover remain unresolved. We performed a field experiment in high and low latitude study sites, with a factorial manipulation of precipitation and herbivory at each site. These factors are hypothesized to be important in driving HCN cline formation, so we investigated their effects on fitness of a white clover F3 recombinant population, segregating for the alleles underlying the HCN chemical defense phenotype. Surprisingly, we found precipitation and herbivory either did not drive differential selection on HCN or its metabolic components, or did not impose selection in a manner consistent with the maintenance of observed HCN clines. Instead, we find that the production only one of the metabolic components of HCN, cyanogenic glycosides, resulted in a fitness advantage, even when lacking the ability to produce HCN. This was most prominent at the northern study site, which is again contrary to expectations. These results suggest additional physiological roles that cyanogenic glycosides may play in adaptation and the evolutionary ecology of white clover. This study highlights the importance of experimental manipulations of environmental factors to understand how selection acts on genes underlying important phenotypic traits, often in unexpected ways.

## Introduction

Spatial and temporal variation in the environment frequently drives patterns of local adaptation and the formation of clines (Endler, 1977; Kawecki and Ebert, 2004; Hereford, 2009). Both biotic and abiotic factors can exhibit variable selection across heterogeneous environments, yet it remains poorly understood how spatial variation in specific environmental factors drives local adaptation and the evolution of clines (Nuismer et al., 2000; Siepielski et al., 2017; Briscoe Runquist et al., 2020). This challenge can be partially resolved by conducting field experiments that manipulate multiple environmental factors that are putative agents of selection acting across spatially separated sites spanning an environmental gradient. These manipulative experiments can serve to further our understanding of how populations adapt to multifaceted environments, with the goal of determining the predictability of adaptation in current and future climates.

Selection imposed by the environment can vary spatially, resulting in genotype-by-environment interactions (G × E), whereby certain genotypes are advantageous under particular biotic and abiotic conditions (Gillespie and Turelli, 1989). However, the relative impacts of specific biotic and abiotic factors on selection are often difficult to discern, highly variable among species, and/or highly dependent on environmental context (Macel et al., 2007; Siepielski et al., 2017; Wright et al., 2018; Briscoe Runquist et al., 2020; Hargreaves et al., 2020). Biotic and abiotic environments also often act concurrently to impose selection and influence evolutionary patterns. A prominent hypothesis that integrates biotic and abiotic selection is the latitudinal herbivory-defense hypothesis (LHDH; Johnson and Rasmann, 2011), which posits that at lower latitudes, warmer and more stable climates support greater abundance and diversity of herbivores, resulting in stronger selective pressure on plants due to herbivory (Coley, 1991; Coley and Barone, 1996; Anstett et al., 2016; Moreira et al., 2018). In this case, a plant functional trait (defense) is simultaneously influenced by multiple G × E interactions, where the environment includes both biotic (herbivory) and abiotic factors (latitude, or by proxy, temperature). However, evidence for and against the LHDH is mixed when examined at a large scale (Moles et al., 2011; Anstett et al., 2016), and indeed the incidence of patterns in support of this hypothesis might depend on the specific biotic and abiotic environmental context in question (Anstett et al., 2014; Moreira et al., 2015; Kooyers et al., 2017; Anstett et al., 2018). These complex results highlight the need for a continued focus on concurrent investigation of biotic and abiotic factors as drivers of spatially variable selection on functional traits along environmental gradients.

The cosmopolitan plant white clover (*Trifolium repens*) offers an ideal study system to investigate environmental factors as drivers of selection on an ecologically important trait. Cyanogenesis, or the production of the antiherbivore defense hydrogen cyanide (HCN), is a chemical defense trait found in over 3000 plant species (Poulton, 1990). The production of HCN in *T. repens* is typically controlled by two Mendelian inherited polymorphic loci (Corkill, 1940; Atwood and Sullivan, 1943; Poulton, 1990). One locus governs the ability to produce cyanogenic glycosides (CNglcs; namely, linamarin and lotaustralin), which is controlled by three linked genes (*CYP79D15*, *CYP736A187*, and *UGT85K17*) in the metabolic pathway that produces linamarin and lotaustralin (Corkill, 1940; Atwood and Sullivan, 1943; Olsen et al., 2008). We hereafter refer to this locus as *Ac*/*ac* (Corkill, 1940; Atwood and Sullivan, 1943; Olsen et al., 2008). The other locus (*Li*/*li*) governs the ability to produce a β-glycosidase called linamarase, a hydrolyzing enzyme that cleaves the sugar moiety of CNglcs from the −C≡N functional group to release toxic HCN (Corkill, 1940; Atwood and Sullivan, 1943; Olsen et al., 2007). Deletion of the functional *Ac* allele results in an inability to produce CNglcs while deletion of the functional *Li* allele results in an inability to produce linamarase (Olsen et al., 2007; Olsen and Small, 2018). Therefore, possession of at least one dominant allele (*Ac* and *Li*) at each locus results in the presence of CNglcs and linamarase, respectively, and the ability to produce HCN. Plants that are homozygous recessive *ac* lack CNglcs, and plants that are homozygous recessive *li* lack linamarase (Corkill, 1940; Atwood and Sullivan, 1943). This genetic architecture results in four cyanogenesis phenotypes (referred to as “cyanotypes”): one cyanotype that can produce both CNglcs and linamarase (HCN^+^/cyanogenic; AcLi); one that can produce only CNglcs (HCN^−^/acyanogenic; Acli); one that can produce only linamarase (HCN^−^; acLi); and one that can produce neither CNglcs nor linamarase (HCN^−^; acli).

Clines in cyanogenesis in *T. repens* have been demonstrated across many types of environmental gradients (Daday, 1954a; Daday, 1954b, Daday, 1958; Ganders, 1990; Thompson et al., 2016). In general, the formation of clines in cyanogenesis relies on shifts in the ratio of benefits to costs of HCN production along environmental gradients, altering selection for or against cyanogenesis. Many hypotheses have been proposed to describe the agents of selection driving cyanogenesis cline formation based on the costs and benefits of HCN production. For example, herbivory and soil moisture have been hypothesized as potential environmental factors driving latitudinal cyanogenesis cline formation based on the benefits of HCN or its metabolic components across environments. Specifically, cyanogenesis may be favoured at lower latitudes due to greater herbivore pressure (Daday, 1965; Angseesing, 1974; Dirzo and Harper, 1982; Kakes, 1989; Santangelo et al., 2019), while CNglcs (and possibly linamarase) could provide additional benefits of nitrogen recycling under low soil moisture conditions, which could vary predictably across the range of white clover (Foulds and Grime, 1972a,b; Kooyers et al., 2014; Albano and Johnson, 2023; Hendrickson et al., 2025). Freezing temperatures have been similarly hypothesized as drivers of latitudinal cyanogenesis cline formation based on exacerbated costs of HCN or its metabolic components in colder environments (Daday, 1965; Olsen and Ungerer, 2008; Kooyers et al., 2018; Fadoul et al., 2023). However, conflicting evidence is present in the literature for each of these hypothesized environmental factors, with common garden studies across latitudes also not definitively uncovering causative selection pressures (Wright et al., 2018; Albano et al., 2024). The importance of multiple biotic and abiotic factors in imposing selection has not yet been directly tested using manipulative experiments in the field. Therefore, to improve our understanding of the mechanisms driving local adaptation and the evolution of observed clines, it is important to perform field experiments across environments that manipulate the effects of multiple environmental factors on a plant species within known genetic control of an ecologically important trait.

To try and resolve uncertainty in the forces driving the evolution of HCN clines in *T. repens*, we first sought to test whether there is a latitudinal cline in herbivory and HCN frequency across an extended latitudinal gradient. We then conducted a manipulative field experiment between disparate latitudes, creating two study sites in the northern and southern distributions of the introduced range of *T. repens* in North America and performing a factorial manipulation of herbivory and precipitation within each site. This experiment tests the overarching hypothesis that biotic and abiotic factors concurrently drive the evolution of adaptive clines in cyanogenesis along latitudinal gradients. Our specific research questions are: (Q1) Is there a cline in herbivory and HCN frequency across a latitudinal gradient in the introduced North American range of *T. repens*? (Q2) Is selection for or against cyanogenesis or its metabolic components influenced by the manipulation of precipitation or herbivory? (Q3) Is selection on cyanogenesis or its metabolic components by precipitation or herbivory also dependent on the latitude of the study site? (Q4) Is the outcome of any observed selection on cyanogenesis or its metabolic components consistent with the maintenance of the cyanogenesis polymorphism within populations or the formation of cyanogenesis clines across latitudinal gradients? We expect to observe strong evidence of latitudinal clines in herbivory and HCN frequency, and selection due to precipitation and herbivory that is consistent with the maintenance of the cyanogenesis polymorphism and the formation of cyanogenesis clines across latitude in North America, based on three specific predictions (**Figure 1**). First, we predict exacerbated costs of producing HCN, CNglcs, and/or linamarase at the higher latitude site due to seasonal freezing temperatures. Second, when herbivores are reduced, we predict a larger beneficial effect on fitness in *T. repens* unable to invest in HCN, CNglcs, and/or linamarase in the lower latitude site, due to preferential investment into growth and reproduction as opposed to defense in plants that are already experimentally defended. Third, we predict *Ac* plants to experience a fitness advantage under drought, and that this advantage could vary between the high and low latitude sites based on differences in the level of precipitation and evapotranspiration occurring throughout the duration of the experiment, with both expected to be higher at the low latitude site, in general. Our results provide insight into the potential interplay between biotic and abiotic environments in the formation of clines in an ecologically important trait within a widespread plant species.

**Figure 1.**
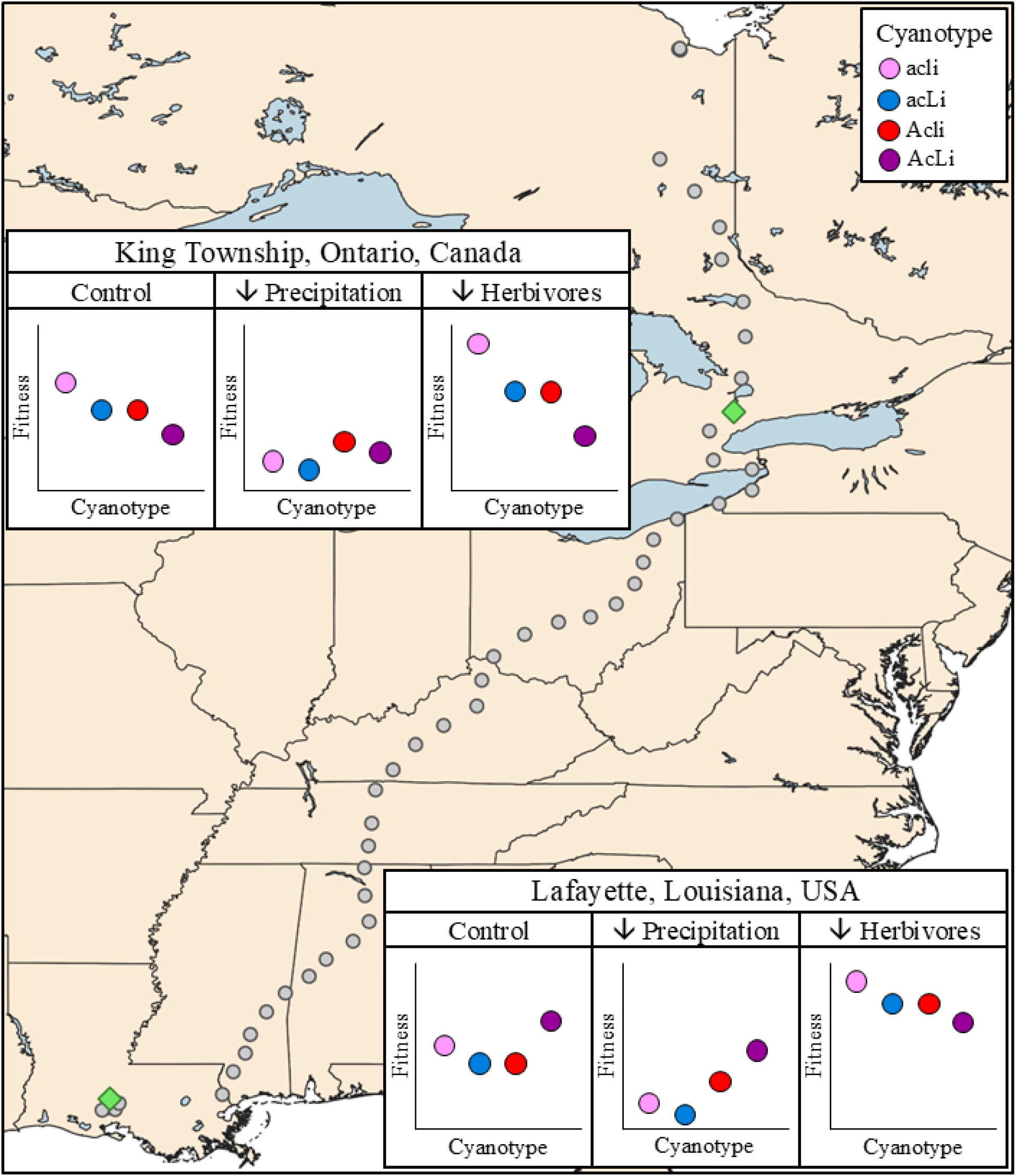
Map identifying study sites, sampled *T. repens* populations, and experiment predictions. The Ontario and Louisiana study sites are identified by green diamonds. The 50 sampled populations used to investigate the presence of a latitudinal cyanogenesis clines in North American *T. repens* are identified by grey circles. Each study site is associated with predictions of relative fitness of *T. repens* cyanotypes under natural conditions and when exposed to precipitation and herbivore manipulations. In Ontario, costs of CNglcs and linamarase are exacerbated in the cooler climate, selecting against Acli, acLi, and AcLi plants under natural/control conditions. Precipitation manipulation reduces fitness to a lesser degree in *Ac* plants due to nitrogen recycling advantages under drought. Herbivore manipulation exacerbates fitness advantages of acyanogenic cyanotypes due to limited metabolic investment into defense compounds when plants are already experimentally defended. In Louisiana, AcLi are selected for under natural conditions due to greater herbivore pressure, while Acli and acLi are selected against due to a lack of defense but increased investment in defense precursors. Precipitation manipulation again reduces fitness to a lesser degree in *Ac* plants due to advantages under drought. Herbivore manipulation again provides relative fitness advantages to acyanogenic cyanotypes due to limited metabolic investment into defense compounds when plants are already experimentally defended, while the relative costs of *Ac* and *Li* are diminished compared to the colder, northern climate.

## Methods

### Study system

*Trifolium repens* is a cosmopolitan perennial legume and obligate outcrosser that can reproduce sexually via flowers and seeds and clonally via stolons, often forming large clonal mats in lawns, fields, and other human-disturbed areas (Burdon, 1983). These multiple modes of reproduction have contributed to the global range expansion of *T. repens* from its native Eurasia to multiple introduced ranges on many other continents, including North America (Daday, 1958; Kjærgaard, 2003). We tested for the presence of latitudinal clines in HCN, the prominent chemical defense mechanism in *T. repens* described above, through field surveys of populations across a large latitudinal transect of the introduced range of North America (**Supplementary Material**)

### Experimental design

All plant material for the experiment is the product of an F3 generation of *T. repens* seeds (methods for experimental plant generation found in **Supplementary Material**; **Figure S1**). We addressed our research questions using experimental manipulations of precipitation and herbivory at two study sites, one located at the Koffler Scientific Reserve <www.ksr.utoronto.ca>, (near King Township, Ontario, Canada; 44°1’40” N, 79°39’29” W) and the other located in Lafayette, Louisiana, USA (30°18’24” N, 92°00’33” W). Historical and current climatic characteristics of each study site can be found in **Table S1**. F3 seedlings were germinated in peat pots (8 cm diameter, 591 mL volume) filled with Promix LP15 (Premier Tech Horticulture, Rivière-du-Loup, Québec, Canada) and established for 8 weeks in a greenhouse near each study site. At the northern study site, plants were established in a climate-controlled greenhouse at 25°C day, 15°C night, and a 14 h:10 h light:dark cycle. At the southern study site, plants were established in a non-climate-controlled greenhouse at ambient conditions. Once cyanotypes were confirmed for each seedling, individuals of each of the four cyanotypes were randomly selected for planting into each of 96 plots, for a total of 384 plants per study site (**Figure S2A**). Each plot was then overlaid with a 1.52 m × 1.82 m sheet of black landscaping fabric, into which four holes were cut to allow for the peat pots containing each individual plant to be embedded into the natural soil at each study site (**Figure S2A**). Individual plants were then randomly assigned and planted into one of the four holes in each plot in spring 2021, with a target of one individual of each cyanotype within each subplot. This target was achieved in Ontario, but in Louisiana, a lack of germination success for some cyanotypes prevented each plot from having exactly one individual of each cyanotype, and seedlings from cyanotypes with an excess number of available individuals were used to ensure four plants were present in each subplot. In total, 50 subplots contained one of each cyanotype, 42 subplots contained no more than two individuals of the same cyanotype, and 4 subplots contained no more than three individuals of the same cyanotype.

In Ontario, our study used a split-plot design, with herbivory manipulated at the whole plot level and precipitation manipulated at the subplot level. The 48 whole plots at each study site were arranged as 96 paired subplots, in which one subplot was assigned as receiving the precipitation reduction (drought) treatment, while the other subplot remained as an ambient precipitation control. A herbivore reduction treatment was then randomly assigned to 24 of the 48 whole plots, resulting in each of the 96 subplots being assigned one of four treatment combinations (**Figure S2A**). In Louisiana, design was similar, but the combination of precipitation reduction and herbivore reduction treatments (four treatment combinations between the two 2-level manipulations) was applied at the subplot level. To manipulate precipitation, rainout shelters were constructed following the guidelines of Kundel et al. (2018), with some modifications based on available materials and experiment requirements. Rainout shelters were constructed with a wooden frame to dimensions of 1.52 m × 1.82 m, to match the dimensions of the landscaping fabric covering each subplot, and were raised in the centre to allow precipitation to run off the subplot (**Figure S2B,C**). Clear, corrugated polycarbonate strips were attached to the frame to cover a defined percentage of the area of the subplot and thereby either act as a precipitation reduction treatment or a control, depending on the orientation in which the polycarbonate strips were attached to the frame (i.e., concave down to allow precipitation onto the plants versus concave up to capture precipitation and carry away from the plants; **Figure S2D**). For the 2021 growing season, an equal number of transparent, corrugated polycarbonate strips (10) were attached to both the precipitation reduction shelters and the control shelters and covered 50% of the subplot area with the intention of excluding ∼50% of precipitation. Soil moisture (percent volume) was measured periodically across each growing season using a soil moisture meter (Delta-T SM150 Soil Moisture Kit and Moisture Sensor; Delta-T Devices, Burwell, UK), with three replicate measurements taken in each subplot in Ontario and four replicate measurements taken in each subplot in Louisiana (**Figure S3**). In year 1 (2021), the rainout shelters did not adequately reduce soil moisture in either Ontario (precipitation reduction mean: 29.2 ± 0.3% volume, control mean: 29.5 ± 0.3% volume; *P* = 0.504) or Louisiana (precipitation reduction mean: 37.7 ± 0.8% volume, control mean: 36.7 ± 0.8% volume; *P* = 0.423; **Table S2**). Therefore, in year 2 (2022), shelters were modified to contain 5 transparent, corrugated polycarbonate strips on the control shelters and 15 strips on the precipitation reduction shelters to cover ∼75% of the area of treated subplots and allowing for a target of excluding ∼75% of precipitation. Additionally, at the Ontario study site only, a drainage pipe was attached to the base of the rainout shelters on the precipitation reduction side of whole plots to catch precipitation and allow it to evaporate. In year 2, soil volumetric water content in precipitation reduction subplots was 18% lower than in control subplots in Ontario (precipitation reduction mean: 29.6 ± 0.2%, control mean: 36.1 ± 0.2%; *P* < 0.001), while rainout shelters still did not adequately reduce soil volumetric water content in Louisiana (precipitation reduction mean: 32.0 ± 1.1%, control mean: 32.4 ± 1.2%; *P* = 0.841; **Table S2**). The years of the experiment were particularly wet in Louisiana, subjecting this study site to flooding during any moderate rain, likely causing the limited effectiveness of the precipitation reduction treatment throughout the experiment (**Table S1**).

To manipulate herbivory, a combination of insecticide and molluscicide was applied biweekly throughout the 2021 and 2022 growing seasons to deter feeding by all common *T. repens* herbivores, following the protocol of Santangelo et al. (2019). Briefly, a 0.0092% solution of Coragen® insecticide (Dupont Chemical, Mississauga, Ontario, Canada) was applied to all aboveground tissue of treated plants until beading of the liquid occurred on leaves, using a pressurized pesticide sprayer, with an equal volume of water being applied to control plants. Additionally, ∼800 mg of Safer’s® Slug and Snail Killer molluscicide pellets (Safer Brand, Lancaster, Pennsylvania, USA) were scattered across the surface of each treated plant. These treatments were shown to be effective at reducing herbivory at both study sites (see **Results**) and therefore were not modified between 2021 and 2022.

All plants were maintained for two full growing seasons at each study site, from spring 2021 until autumn 2022, with scheduled measurements of plant fitness being conducted across each growing season. Whether a plant survived and/or flowered was assessed weekly, with ripe fruits also being counted and collected weekly. Seeds were then separated using a thresher (Precision Machine Co., Lincoln, NE, USA), and fecundity was measured as total seed set mass per plant. Photos were taken of each plant monthly and plant area was determined using Easy Leaf Area (Version 2.0; Easlon and Bloom, 2014). The maximum plant area across each growing season was determined and growth rate was calculated as (ln[A_max_] – ln[A_1_])/(t_max_ – t_1_), where A_max_ represents the maximum plant area, A_1_ represents the initial plant area (i.e., after one month of growth in the field), and (t_max_ – t_1_) represents the number of days between the photos taken for maximum plant area and initial plant area. Herbivory was assessed visually to the nearest 1% from 5 fully-expanded trifoliate leaves per plant (or fewer if a plant had less than 5 fully-expanded leaves) each month following the methods of Johnson et al. (2016). In Louisiana, aboveground plant tissue outside of an 8 cm diameter area around the planting location of each individual was manually removed in January 2022 to prevent plants from growing too large and competing with neighbouring individuals during the second growing season. In Ontario, winter freezing temperatures result in a natural dieback of vegetation after each growing season, so manual tissue removal was not necessary.

### Statistical analysis

All statistical analyses were conducted in R Version 4.4.1 (2024) and R Studio Version 2024.09.1 (R Development Core Team, 2024). All response variables were then analyzed through general linear mixed effects models using the function *glmmTMB* in the *glmmTMB* package (Version 1.1.9; Brooks et al., 2017), with χ^2^ and *p* values calculated using the *Anova* function in the *car* package (Version 3.1-2; Fox and Weisberg, 2019) with Type II sums-of-squares. For significant main effects of cyanotype and any significant interactions of cyanotype with one or more of the manipulated variables, custom contrasts were created using the *emmeans* and *contrast* functions in the *emmeans* package (Version 1.10.4; Lenth, 2024). For models using quantitative measures of fitness or herbivory, all transformations necessary to meet assumptions of normality and homogeneity of variance can be found in **Table S3**.

Each of the response variables (survival, whether plants produced flowers, whether plants produced seeds, number of flower heads, seed set mass, maximum plant area, growth rate, and herbivory) were assessed to determine the effects of cyanotype and each of our experimental manipulations on plant fitness. The number of flower heads and seed set mass produced by each plant were both zero-inflated, so prior to analyzing these variables, it was necessary to first analyze whether plants flowered or produced seeds, and then subset the number of flower heads and seed set mass variables to only include values greater than zero, as has been done previously (Anderson et al., 2015; Ferris and Willis, 2018; Albano and Johnson, 2023; Scharnagl et al., 2023). In Ontario, all models consisted of the following structure:

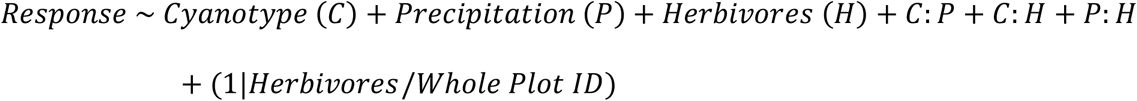

In Louisiana, all models consisted of the following structure:

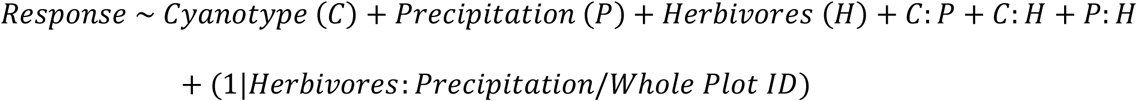

The split-plot experimental design necessitated the use of nested random effects to account for non-independence of replicates within each whole plot in Ontario, while the different treatment application in Louisiana necessitated an interaction term between the herbivore and precipitation reduction treatments as part of the structure of random effects. Models including additional nested random effects to account for non-independence of replicates within each subplot either experienced convergence issues or did not alter our conclusions and were therefore not used. Cyanotype (AcLi, Acli, acLi, or acli), precipitation manipulation (reduced versus control) and herbivore manipulation (reduced versus control) were included as fixed effects along with all two-way interactions. The three-way interaction was also originally included in all models but was not statistically significant at *p* < 0.05 in any cases and therefore was removed from all models to increase statistical power in testing the remaining fixed effects. Data from each of the two growing seasons were analyzed separately due to the difference in how the precipitation manipulation was conducted between the 2021 and 2022 growing seasons and the manual removal of aboveground tissue in Louisiana after the 2021 growing season. Data from each of the two study sites were analyzed separately due to some subplots not containing exactly one individual of each cyanotype in Louisiana, along with the differences in treatment application.

## Results

### Field surveys

Our field surveys of *T. repens* populations identified the presence of clines in both herbivory and plant defense across a north-south temperature gradient in North America. Mean annual temperature was strongly negatively correlated with latitude (r^2^ = 0.995; *P* < 0.001), with higher latitudes being associated with lower mean annual temperatures (**Figure 2A**). The proportion of cyanogenic *T. repens* plants was strongly positively correlated with mean annual temperature (χ^2^ = 21.624; *P* < 0.001), with warmer climates (i.e., lower latitudes) being associated with greater incidence of cyanogenesis (**Figure 2B**). The percent of leaf area consumed was weakly positively correlated with mean annual temperature (r^2^ = 0.065; *P* = 0.075; **Figure 2C**), with warmer climates (i.e., lower latitudes) being associated with marginally higher levels of herbivory.

**Figure 2.**
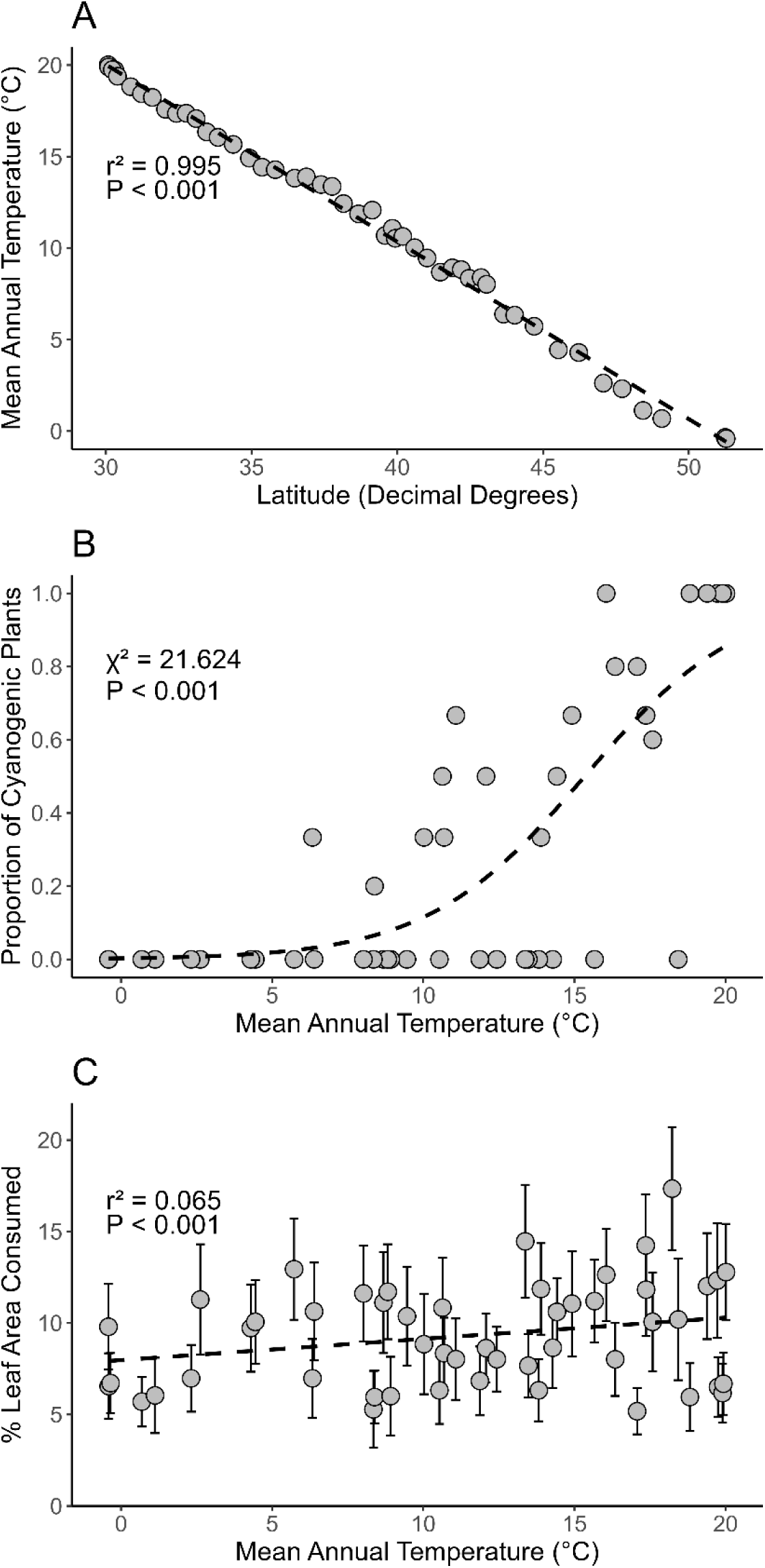
Clines in cyanogenesis and herbivory in the North American range of *T. repens* (A) The relationship between latitude and mean annual temperature for 50 sampling locations across a north-south transect in North America. (B) The effect of mean annual temperature on the proportion of cyanogenic *T. repens* individuals (N = 140) from 47 of the sampling locations. (C) The effect of mean annual temperature on the mean percentage of leaf area consumed for 50 individuals (1 trifoliate leaf per individual) assessed at each of the 50 sampled locations.

### Plant sexual fitness in Year 1 (2021)

In Ontario, cyanotype affected plant fitness, but experimental manipulations (precipitation and herbivores) did not affect selection on *T. repens* cyanotypes (**Table 1**). Specifically, cyanogenic plants had 54% larger seed set mass than acyanogenic plants and *Ac* (CNglc-producing) plants had 56% larger seed set mass than *ac* plants, while *Li/li* did not affect seed set mass, and cyanotype did not affect whether plants produced seeds (**Figure 3**). We found limited evidence of an effect of the precipitation manipulation on the sexual fitness of *T. repens*, potentially due to the lack of effectiveness of this treatment in 2021. Similarly, the herbivore manipulation did not affect the sexual fitness of *T. repens* in Ontario in 2021, even though it reduced herbivory by 54%.

**Figure 3.**
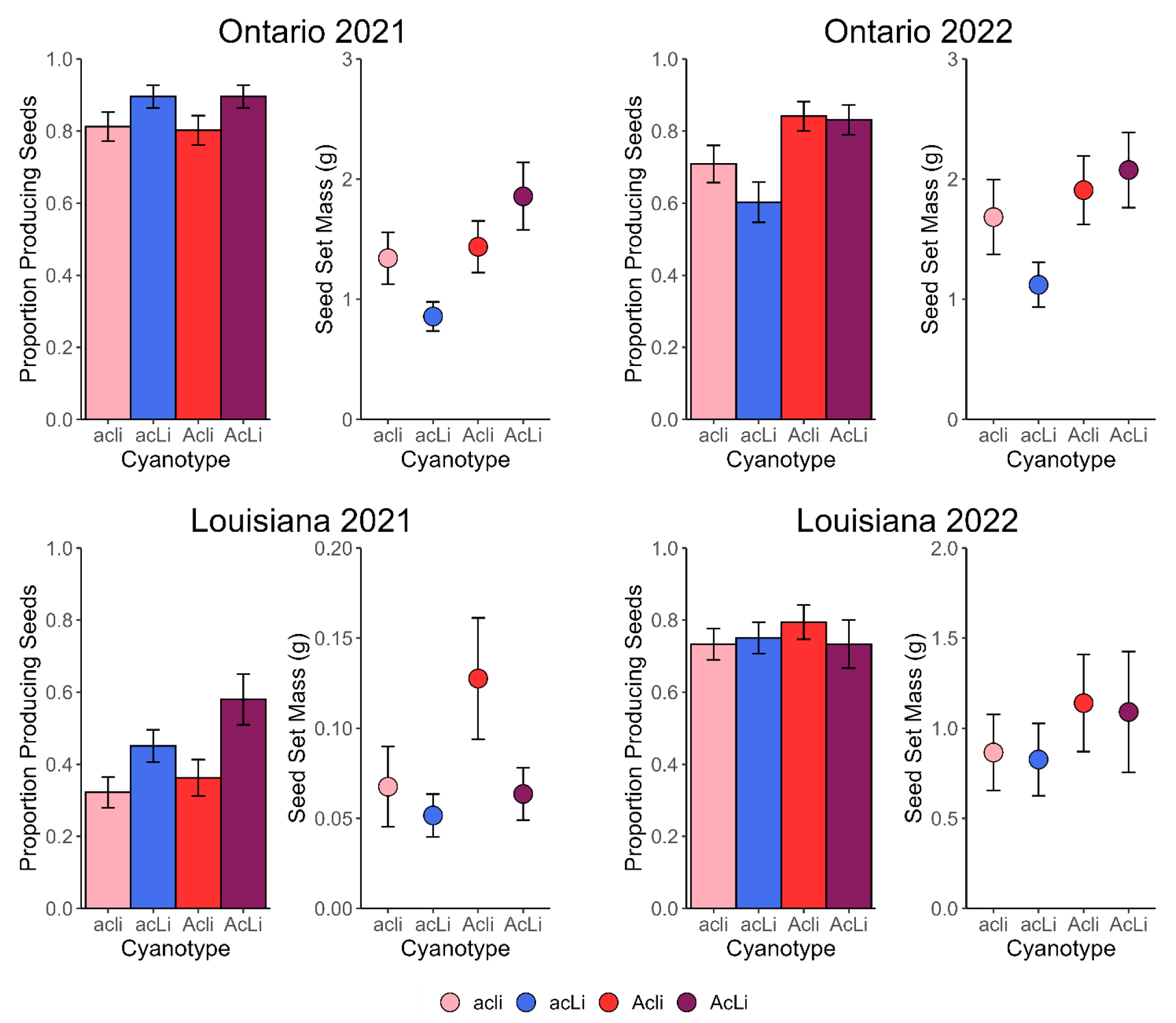
The effect of cyanotype on plant fitness. We show the proportion of *T. repens* plants of each cyanotype (acli, acLi, Acli, AcLi) that produced seeds and their total seed mass in Ontario in 2021 (top left) and 2022 (top right) and in Louisiana in 2021 (bottom left) and 2022 (bottom right). Error bars are ± one standard error for each mean value.

**Table 1.** Results of mixed effects models assessing how cyanotype, manipulation of precipitation, manipulation of herbivore pressure, and their two-way interactions affected the proportion of *T. repens* plants producing seeds and the seed set mass in Ontario and Louisiana study sites in the 2021 and 2022 growing seasons. Custom contrasts were performed for significant main effects of cyanotype to test the specific effects of cyanogenesis (AcLi vs. Acli + acLi + acli), presence of the dominant *Ac* allele (AcLi + Acli vs. acLi + acli), and presence of the dominant *Li* allele (AcLi + acLi vs. Acli + acli) on the fitness of *T. repens*. Signs of > and < are with respect to the first listed side of the custom contrast (cyanogenic, *Ac*, or *Li*). Significant effects (*P* < 0.05) are listed in bold.

| Ontario Study Site | 2021 |  |  |  |  |  | 2022 |  |  |  |  |  |
| --- | --- | --- | --- | --- | --- | --- | --- | --- | --- | --- | --- | --- |
|  | Prop. Producing Seeds |  |  | Seed Set Mass |  |  | Prop. Producing Seeds |  |  | Seed Set Mass |  |  |
| Predictor | $\chi^2$ | df | *P* | $\chi^2$ | df | *P* | $\chi^2$ | df | *P* | $\chi^2$ | df | *P* |
| Cyanotype (C) | 7.431 | 3 | 0.059 | **16.516** | **3** | **0.001** | **15.635** | **3** | **0.001** | **9.024** | **3** | **0.029** |
| Precipitation (P) | 0.051 | 1 | 0.822 | 0.266 | 1 | 0.606 | 0.893 | 1 | 0.345 | **6.070** | **1** | **0.014** |
| Herbivores (H) | 1.642 | 1 | 0.200 | 0.000 | 1 | 0.992 | **4.507** | **1** | **0.034** | 2.811 | 1 | 0.094 |
| C × P | 5.644 | 3 | 0.130 | 4.414 | 3 | 0.220 | **8.456** | **3** | **0.037** | 3.348 | 3 | 0.341 |
| C × H | 2.275 | 3 | 0.517 | 1.342 | 3 | 0.719 | 2.743 | 3 | 0.433 | 0.204 | 3 | 0.977 |
| P × H | 0.001 | 1 | 0.974 | 0.924 | 1 | 0.336 | 2.873 | 1 | 0.090 | 0.650 | 1 | 0.420 |
| Contrast | Sign |  | *P* | Sign |  | *P* | Sign |  | *P* | Sign |  | *P* |
| *Cyanotype* |  |  |  |  |  |  |  |  |  |  |  |  |
| Cyanogenic vs. Acyanogenic | — |  | — | > |  | **0.009** | = |  | 0.062 | = |  | 0.057 |
| *Ac* vs. *ac* | — |  | — | > |  | **0.002** | > |  | **< 0.001** | > |  | **0.026** |
| *Li* vs. *li* | — |  | — | = |  | 0.403 | = |  | 0.170 | = |  | 0.393 |
| Louisiana Study Site | 2021 |  |  |  |  |  | 2022 |  |  |  |  |  |
|  | Prop. Producing Seeds |  |  | Seed Set Mass |  |  | Prop. Producing Seeds |  |  | Seed Set Mass |  |  |
| Predictor | $\chi^2$ | df | *P* | $\chi^2$ | df | *P* | $\chi^2$ | df | *P* | $\chi^2$ | df | *P* |
| Cyanotype (C) | **11.694** | **3** | **0.009** | 1.345 | 3 | 0.719 | 0.919 | 3 | 0.821 | **10.522** | **3** | **0.015** |
| Precipitation (P) | 2.386 | 1 | 0.122 | 0.074 | 1 | 0.786 | 0.790 | 1 | 0.374 | 0.155 | 1 | 0.694 |
| Herbivores (H) | **25.264** | **1** | **< 0.001** | **13.484** | **1** | **< 0.001** | 0.352 | 1 | 0.553 | 0.338 | 1 | 0.561 |
| C × P | 4.468 | 3 | 0.215 | 0.291 | 3 | 0.962 | 1.503 | 3 | 0.682 | 0.717 | 3 | 0.869 |
| C × H | 1.785 | 3 | 0.618 | **10.453** | **3** | **0.015** | 0.088 | 3 | 0.993 | 1.370 | 3 | 0.713 |
| P × H | 0.574 | 1 | 0.449 | 0.333 | 1 | 0.564 | 2.449 | 1 | 0.118 | 0.480 | 1 | 0.488 |
| Contrast | Sign |  | *P* | Sign |  | *P* | Sign |  | *P* | Sign |  | *P* |
| *Cyanotype* |  |  |  |  |  |  |  |  |  |  |  |  |
| Cyanogenic vs. Acyanogenic | > |  | **0.009** | — |  | — | — |  | — | = |  | 0.893 |
| *Ac* vs. *ac* | = |  | 0.114 | — |  | — | — |  | — | > |  | **0.023** |
| *Li* vs. *li* | > |  | **0.001** | — |  | — | — |  | — | = |  | 0.125 |

In Louisiana, cyanotype affected whether plants produced seeds but not their seed set mass, while the herbivore manipulation affected plant fitness and selection on *T. repens* cyanotypes (**Table 1**; **Figure 3**). Cyanogenic (HCN-producing) plants were 53% more likely to produce seeds than acyanogenic plants and *Li* (linamarase-producing) plants were 44% more likely to produce seeds than *li* plants, while *Ac*/*ac* did not affect whether plants produced seeds (**Figure 3**). We found strong evidence of an effect of the herbivore manipulation on the sexual fitness of *T. repens*, with herbivore reduction resulting in 2.5× more individuals producing seeds and 3.2× greater seed set mass. Cyanotype and herbivore manipulation also interacted to affect the sexual fitness of *T. repens*, with AcLi, Acli, and acli plants producing 4.5×, 11.2×, and 5.8× larger seed set masses, respectively, when herbivores were reduced versus the control, while acLi plants exhibited no sexual fitness advantage when herbivores were reduced (**Figure 4**). The precipitation manipulation had no effect on fitness or selection, again likely due to the lack of effectiveness of this experimental treatment in 2021.

**Figure 4.**
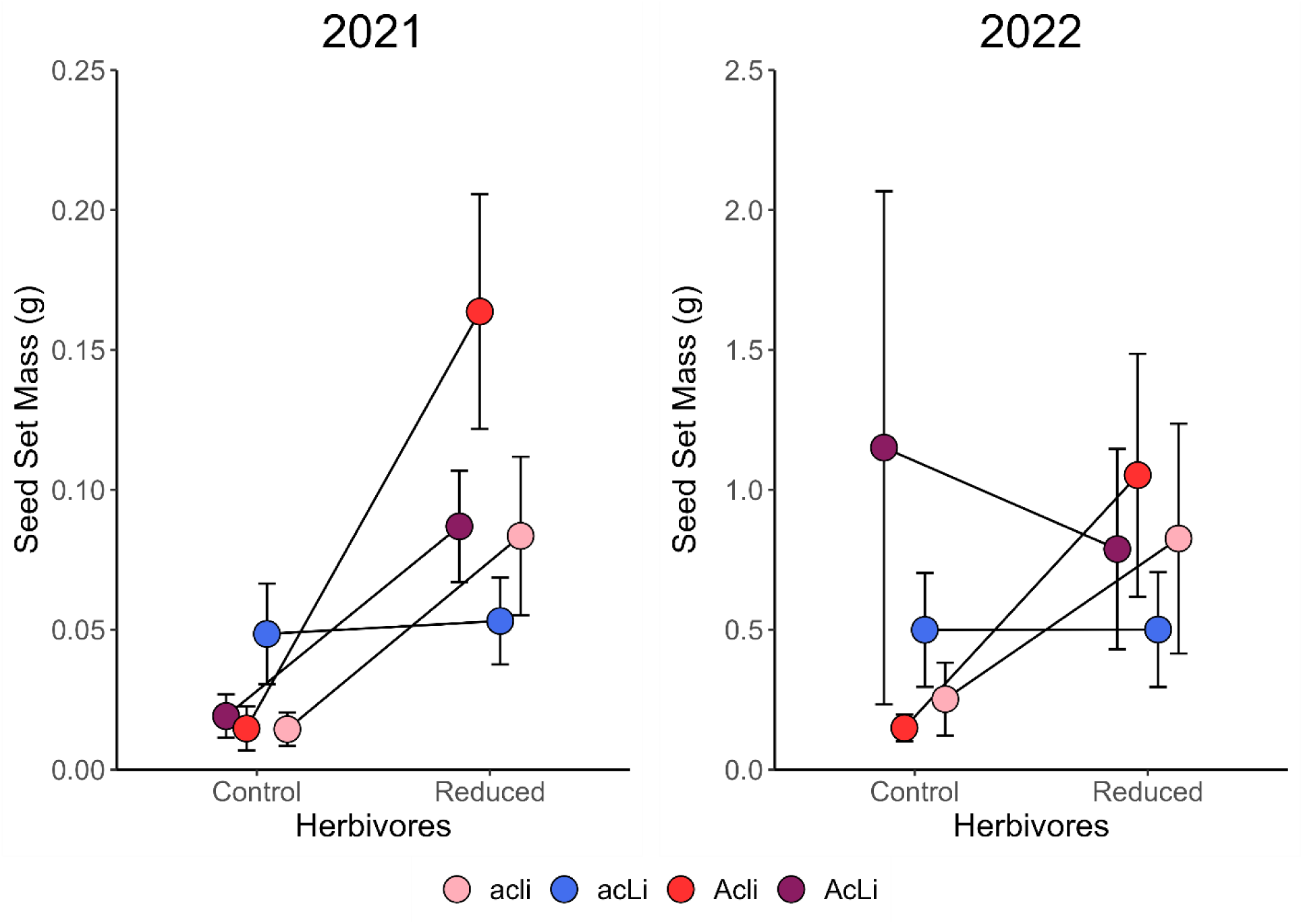
Herbivore manipulation imposes differential selection on cyanotypes in Louisiana in 2021. We show the effect of cyanotype (acli, acLi, Acli, AcLi) and herbivore manipulation (control vs. reduced) on seed set mass produced by *T. repens* plants at the Louisiana study site in 2021 and 2022. Error bars are ± one standard error for each mean value.

### Plant sexual fitness in Year 2 (2022)

In Ontario, cyanotype affected plant fitness, but we only found limited evidence of the precipitation manipulation affecting selection on *T. repens* cyanotypes and no evidence of an effect of the herbivore manipulation on selection (**Table 1**). Specifically, cyanogenic plants were 15% more likely to produce seeds than acyanogenic plants and *Ac* plants were 27% more likely to produce seeds than *ac* plants (**Figure 3**). Furthermore, cyanogenic plants produced 27% greater seed set mass than acyanogenic plants and *Ac* plants produced 41% greater seed set mass than *ac* plants. *Li/li* did not affect seed set mass or whether plants produced seeds (**Figure 3**). Precipitation reduction resulted in 36% smaller seed set mass, while cyanotype and precipitation manipulation also interacted to affect whether plants produced seeds (**Table 1**), with acli plants 54% more likely to produce seeds under reduced precipitation, while AcLi, Acli, and acLi plants were not affected by precipitation reduction (**Figure 5**). Herbivore reduction resulted in a 21% higher proportion of individuals producing seeds, but it did not affect seed set mass (**Table 1**).

**Figure 5.**
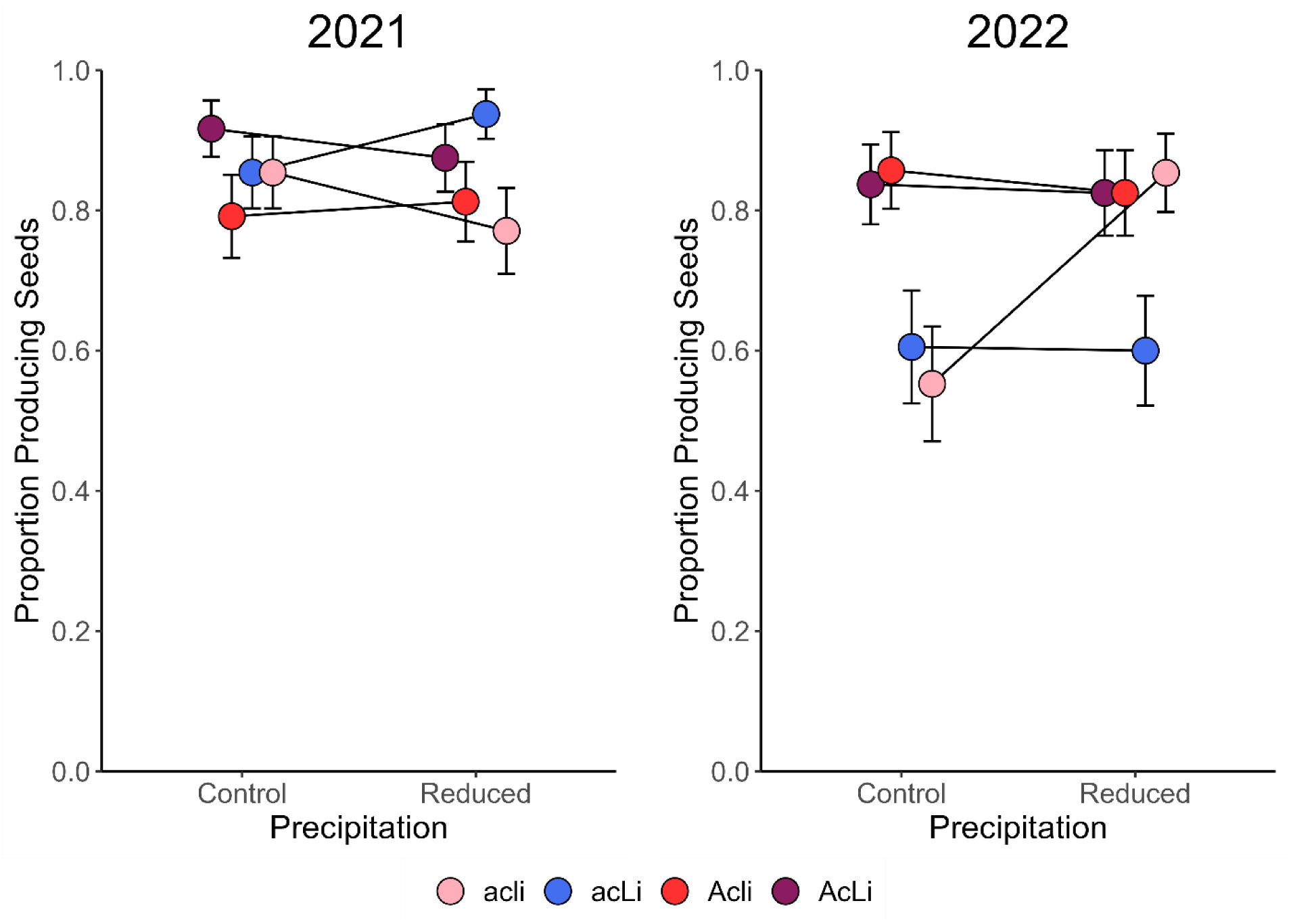
Precipitation manipulation imposes differential selection on cyanotypes in Ontario in 2022. We show the effect of cyanotype (acli, acLi, Acli, AcLi) and precipitation manipulation (control vs. reduced) on the proportion of *T. repens* plants that produced seeds at the Ontario study site in 2021 and 2022. Error bars are ± one standard error for each mean value.

In Louisiana, cyanotype affected plant fitness, but experimental manipulations (precipitation and herbivores) did not affect selection on *T. repens* cyanotypes (**Table 1**). Specifically, *Ac* (CNglc-producing) plants had 32% larger seed set mass than *ac* plants, while cyanogenesis and *Li/li* did not affect seed set mass, and cyanotype did not affect whether plants produced seeds (**Figure 3**). We again found limited evidence of an effect of the precipitation manipulation on the sexual fitness of *T. repens*, despite the rainout shelter modifications made prior to the 2022 growing season. Similarly, the herbivore manipulation did not affect the sexual fitness of *T. repens* in Louisiana in 2022, even though it reduced herbivory by 17%.

### Additional reproductive and growth traits in Years 1 and 2

In general, results for whether plants flowered and the number of flower heads produced were qualitatively similar to whether plants produced seeds and seed set mass across both growing seasons at both the Ontario (**Tables S4**, **S6**) and the Louisiana (**Tables S5**, **S7**) study sites. We also did not find evidence of selection for or against *T. repens* cyanotypes due to either precipitation or herbivore manipulation based on measures of survival or vegetative fitness (maximum plant area or growth rate; **Tables S4-S7**).

### Herbivory in Years 1 and 2

In Ontario in 2021, herbivory (measured as the percentage of leaf area consumed) was, as expected, strongly affected by the reduction of herbivores, which resulted in 57% less leaf tissue consumed (herbivore reduction mean: 6.0 ± 0.5% leaf area consumed, control mean: 14.0 ± 0.8% leaf area consumed; **Table S4**). Cyanogenic plants experienced 33% less herbivory than acyanogenic plants (**Figure 6**). Surprisingly, acyanogenic *Ac* (Acli) plants experienced 46% less herbivory than *ac* plants, which is contrary to expectations because solely producing CNglcs does not result in *T. repens* plants being able to produce HCN as a defense mechanism (**Figure 6**). Similarly, in Ontario in 2022, herbivory was strongly affected by the reduction of herbivores, which resulted in 57% less leaf tissue consumed (herbivore reduction mean: 10.0 ± 0.7% leaf area consumed, control mean: 17.8 ± 1.0% leaf area consumed; **Table S6**). Cyanogenic plants experienced 25% less herbivory than acyanogenic plants (**Figure 6**). Acyanogenic *Ac* (Acli) plants experienced 27% less herbivory than *ac* plants, which again, is contrary to expectations (**Figure 6**).

**Figure 6.**
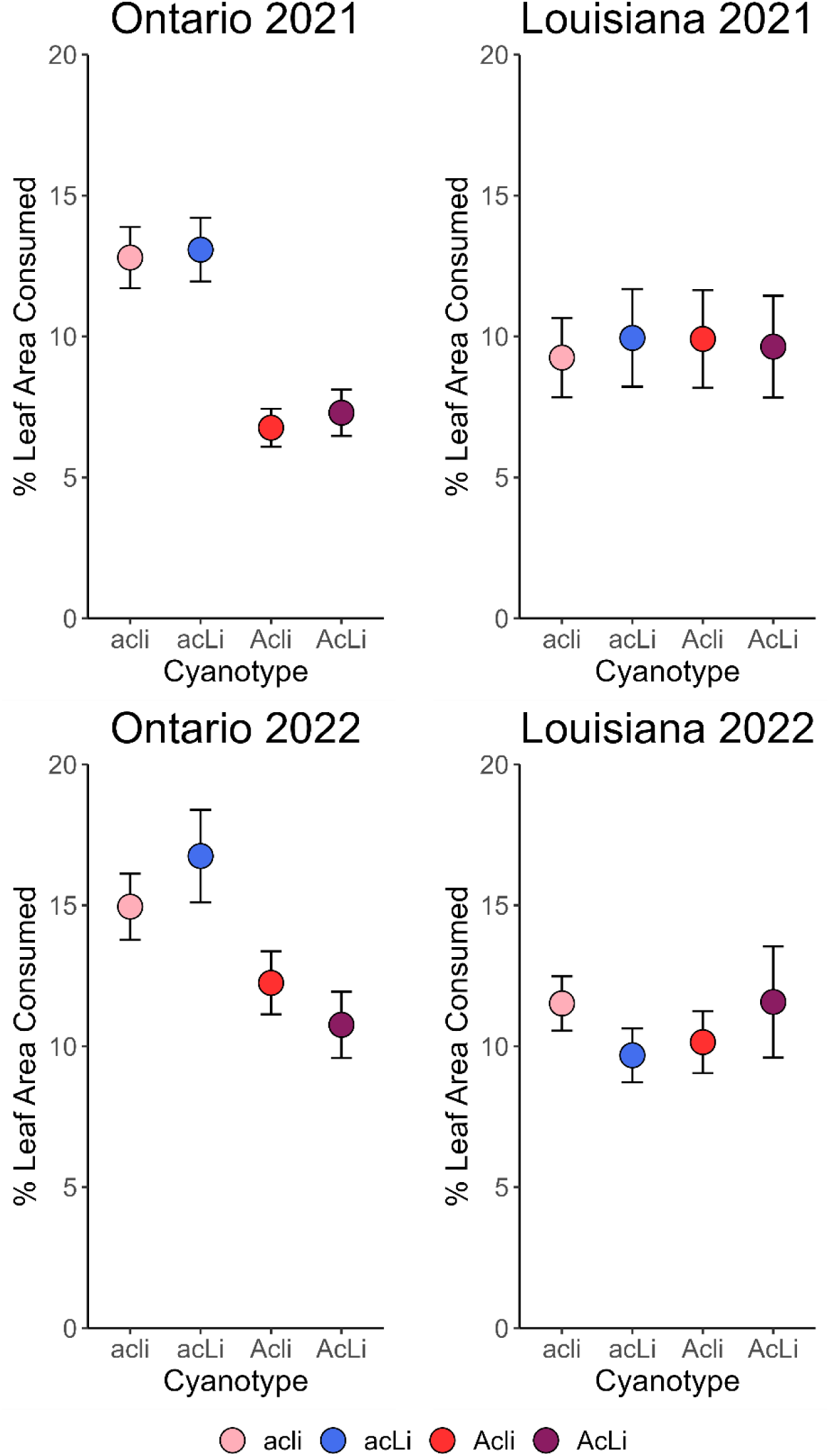
The effect of cyanotype on herbivory. We show the percent of leaf area consumed for *T. repens* cyanotypes at the Ontario (acli, acLi, Acli, AcLi) study site (top left) and the Louisiana study site (top right) in the 2021 growing season and at the Ontario study site (bottom left) and the Louisiana study site (bottom right) in the 2022 growing season. Error bars are ± one standard error for each mean value.

In Louisiana in 2021, herbivory was strongly affected by the reduction of herbivores, which resulted in 87% less leaf area consumed (herbivore reduction mean: 2.2 ± 0.3% leaf area consumed, control mean: 16.6 ± 1.3% leaf area consumed; **Table S5**). However, cyanotype did not affect the amount of herbivory on *T. repens* plants, indicating a lack of effectiveness of HCN in defending plant tissue at this site (**Figure 6**). In Louisiana in 2022, neither the herbivore manipulation nor cyanotype alone affected the amount of herbivory on *T. repens* plants (**Table S7**; **Figure 6**). However, cyanotype and the herbivore manipulation did interact to affect herbivory, with AcLi, Acli, and acLi plants experiencing 47%, 29%, and 23% reductions in leaf area consumed, respectively, when herbivores were reduced versus the control, while acli plants exhibited 15% greater herbivory when herbivores were reduced (**Table S7**).

## Discussion

Our findings indicate the presence of a latitudinal cline in plant defense, with a reduced frequency of cyanogenic *T. repens* plants in colder North American climates, in accordance with the LHDH. Additionally, we provide the first direct evidence of a positive relationship between mean annual temperature and herbivory in *T. repens*, indicating greater herbivore pressure at lower latitudes, despite greater incidence of cyanogenesis, also in accordance with the LHDH (Q1). Our manipulative experiment clearly shows that the metabolic components of HCN are under natural selection, but our results do not support the long-standing hypotheses that precipitation and herbivory impose selection on the presence/absence of HCN or its metabolic components in *T. repens* and drive the formation of cyanogenesis clines. Despite clear evidence that reduced precipitation and herbivory both decrease *T. repens* fitness, we found only limited evidence of precipitation and herbivory imposing differential selection on *T. repens* cyanotypes (Q2). We did find some evidence of differential effects of cyanotype that depended on the study site location, with greater beneficial effects of cyanogenesis and the ability to produce CNglcs on plant fitness at the higher latitude study site, which is contrary to expectations (Q3). In the limited instances when herbivory or precipitation did interact to affect *T. repens* fitness, the outcomes were also contrary to expectations, with demonstrated fitness advantages for unexpected cyanotypes based on the observed latitudinal cyanogenesis cline (Q4). Here we discuss our results and their implications in the context of plant-herbivore interactions and latitudinal cline formation for the presence of HCN and its metabolic components in *T. repens*.

### Limited evidence of precipitation as a driver of cyanogenesis clines

The ability of plants to invest in essential functions like vegetative growth, reproduction, and defense is dependent on the costs and benefits of those investments. Soil moisture availability is a key environmental factor determining the balance of costs and benefits of these functions by impacting resource availability and therefore, the ability of plants to invest into both primary and secondary metabolic processes (Coley et al., 1985; Mundim and Pringle, 2018). Unexpectedly, precipitation had inconsistent effects on *T. repens* fitness. This may be at least partially due to the design of our rainout shelters, which only reduced precipitation in Ontario in 2022, and/or due to consistent flooding that occurred throughout the entire experiment in Louisiana. We also recognize the importance of extreme events in driving natural selection (Grant et al., 2017), with variation in climatic and experimental conditions between years in our experiment possibly explaining observed inconsistencies (**Table S1**, **Table S2**). Nonetheless, we can conclude that even in instances where precipitation did exhibit strong effects on quantitative measures of plant fitness (e.g., precipitation reduction resulted in decreased seed set mass in Ontario in 2022), there were no differential effects on fitness of *T. repens* cyanotypes due to the precipitation manipulation. Thus, plants with different capacities to invest in plant defense did not differ in their ability to invest in growth or reproduction based on the impact of precipitation reduction on resource acquisition, which is contrary to hypothesis surrounding the involvement of water availability in cyanogenesis cline formation. More specifically, our findings are contrary to recent evidence that *Ac* plants exhibit a fitness advantage when experiencing moderate and consistent drought conditions (Kooyers et al., 2014; Albano and Johnson, 2023), potentially due to the ability of plants to recycle N-rich compounds such as CNglcs during periods of low soil moisture (see further discussion below; Møller, 2010; Machingura et al., 2016). Our results are partially supported, however, by classic work on the effects of soil moisture on *T. repens* cyanotypes, with *ac* plants exhibiting greater survival and similar dry biomass compared to *Ac* plants experiencing drought, possibly due to linkage of *Ac*/*ac* with other loci involved in environmental responses (Foulds and Grime, 1972a). As a whole, we are unable to conclude that additional benefits aside from defense, such as greater facilitation of N recycling under drought by *Ac* plants, are involved in cyanogenesis cline formation in *T. repens*.

In our study, a sole instance of differential effects of precipitation reduction on *T. repens* cyanotypes (at the Ontario study site in 2022) resulted in increased likelihood of flowering and producing seeds in acli plants, but not in any other cyanotype. However, when precipitation was reduced, acli plants were still not more likely than Acli or AcLi plants to flower or produce seeds, which would be expected if acyanogenic plants are selected for in colder climates.

Furthermore, our results provide little evidence of selection on HCN, CNglcs, or linamarase due to soil moisture in terms of quantitative measures of fitness (i.e., number of flower heads or seed set mass). The literature remains conflicted on how soil moisture is likely to influence selection on cyanogenesis or its metabolic components in *T. repens*. Foulds and Grime (1972a,b) found a lack of flowering success for cyanogenic plants under drought and a lower frequency of *Ac* in droughted environments, while Kooyers et al. (2014) found a higher frequency of *Ac* in droughted environments. None of these previous studies found an effect of soil moisture availability on the frequency of *Li* or HCN in *T. repens* populations. Overall, our results are inconsistent with the maintenance of cyanogenesis clines across latitudinal gradients and we are unable to provide more definitive insight into the conflicting evidence from the literature concerning the ability of soil moisture/precipitation to drive the formation of *T. repens* cyanogenesis clines. Our methods have improved on previous experiments by manipulating *Ac* and *Li* to be on a randomized genetic background to minimize the influence of other genes involved in environmental response that may have affected previous results. However, because our F3 generation is the product of an original cross of only two parent plants and only a single generation of random crossing, it is possible that loci that are very tightly linked to *Ac*/*ac* or *Li*/*li* are influencing our results as well (Kuo et al., 2024a). We are also unable to account for copy number variation at either locus in the parent plants or among the F3 generation individuals used in the experiment, which could affect rapid adaptation across environmental gradients (Kuo et al., 2024b). A genome-wide association analysis could help elucidate if nearby genomic regions or copy number variation at cyanogenesis loci influence drought response in *T. repens*.

### Costs and benefits of cyanogenesis and its metabolic components as a chemical defense mechanism

The benefits associated with plant chemical defense production are likely to be dependent on environmental context. For example, under the LHDH, the benefits of antiherbivore defense are predicted to be more prominent at lower latitudes due to greater herbivore pressure imposing stronger selection in these environments (Coley, 1991; Coley and Barone, 1996; Johnson and Rasmann, 2011). In line with the LHDH, we demonstrated the presence of a latitudinal cline in herbivory that coincides with clines in the frequency of cyanogenesis in *T. repens* populations, with greater herbivory and cyanogenesis frequency being associated with warmer temperatures (i.e., lower latitudes). If the LHDH is further supported by the results of our manipulative experiment, in Louisiana (where experimental release of herbivore pressure is hypothesized to be most substantial), we would expect interactions between cyanotype and herbivore manipulation, with the reduction of herbivores resulting in the greatest benefit for plants that invest the least amount of resources into defense, and vice versa. Instead, any demonstrated interactions between cyanotype and herbivore manipulation in Louisiana were not concordant with expectations, because acli plants did not benefit the most from the experimental exclusion of herbivores, nor did AcLi plants benefit the least (**Figure 4**).

Our results provide stronger evidence of the presence of CNglcs providing a fitness advantage to *T. repens* than the presence of HCN itself. This is an unexpected result because without the ability to produce linamarase, *Ac* plants are unable to endogenously produce HCN at the moment that cell lysis occurs when herbivores damage plant tissue, which is the typical mechanism by which cyanogenic (AcLi) plants exhibit feeding deterrence. However, Acli plants experienced equal deterrence of herbivory to AcLi plants in Ontario, despite a lack of HCN production. Previous work does indicate the potential for CNglc-producing plants to be able to elicit a toxic effect on herbivores later in the feeding process that could result in deterrence of feeding (Dirzo and Harper, 1982; Kakes, 1989; Desroches et al., 1997; Tattersall et al., 2001; Pankoke et al., 2012). This mechanism is a result of β-glycosidases produced by herbivores, some of which may unintentionally hydrolyze CNglcs to produce HCN in the herbivore’s gut. In fact, previous studies involving feeding assays on *T. repens* tissue indicate the potential for feeding deterrence by CNglc-producing plants (Dirzo and Harper, 1982; Kakes 1989). Dirzo and Harper (1982) also determined that excised guts of slug species frequently produce HCN when exposed to chemical CNglcs. There is evidence for this downstream antiherbivore effect in numerous plant study systems that are able to produce various CNglcs as well, including vicine in broad beans (Desroches et al., 1997), iridoid glycosides in *Plantago lanceolata* (Pankoke et al., 2012), and dhurrin in *Arabidopsis thaliana* and other plant species (Tattersall et al., 2001; Yadav et al., 2023). However, previous feeding trials using *T. repens* also provide evidence of a lack of feeding deterrence in CNglc-producing plants (Dirzo and Harper, 1982; Fadoul et al., 2023), indicating a level of heterogeneity either within or among herbivore species in their production of β-glycosidases that could cause HCN production in the gut post-feeding. A future study conducting a direct test of this mechanism on multiple herbivore communities in natural settings is critical to assessing herbivory as a potential hypothesis to describe the consistent presence of latitudinal clines in HCN (and in the frequency of *Ac*) in *T. repens* populations.

While CNglcs may be providing a fitness advantage to *T. repens* plants through a reduction of herbivory despite a lack of HCN production, it is unlikely that CNglcs acting as a chemical defense mechanism is their only benefit to plant performance. In the sole instance of herbivore manipulation imposing differential effects on the fitness of *T. repens* cyanotypes (at the Louisiana study site in 2021), the reduction of herbivores resulted in the greatest benefit for Acli plants, indicating a distinct selective advantage of CNglc production that outweighs its cost, even when herbivory is experimentally reduced by over 50%. It is possible that the recycling of CNglcs into usable N through β-cyanoalanine synthase serves as this additional benefit to CNglc production (Machingura et al., 2016). The ability to produce CNglcs has been shown to be particularly advantageous under low soil moisture in a greenhouse (Albano and Johnson, 2023), which was not the case in the current study conducted in natural conditions. However, it remains a possibility that this additional source of N is universally beneficial to a plant’s ability to invest in growth and/or reproduction, regardless of soil moisture conditions. By contrast, we did not find consistent evidence of the ability to produce linamarase providing a fitness benefit to *T. repens* plants aside from the contribution of linamarase to HCN production in AcLi plants. Thus, linamarase may incur an allocation cost to plants in the absence of HCN and herbivores. Further work directly addressing all potential costs and benefits of CNglcs and linamarase is needed to more definitively test hypotheses surrounding the formation of cyanogenesis clines in *T. repens*.

### Maintenance of cyanogenesis polymorphisms and clines

We find evidence of epistatic interactions between the *Ac*/*ac* and *Li*/*li* loci but not in a manner that explains the maintenance of the cyanogenesis polymorphism within populations. One such epistatic interaction is the increase in flowering success of acli plants under drought that is not present in any other cyanotype, indicating an absence of both metabolic components of cyanogenesis is required for drought to affect flowering in *T. repens*. An additional epistatic interaction is evidenced by fitness benefits (and greater relative deterrence of herbivory) for cyanogenic plants in Ontario but not in Louisiana, indicating the ability to produce both metabolic components of cyanogenesis provides fitness advantages that are variable among *T. repens* populations. However, these results conflict with expected trends that would explain the maintenance of cyanogenesis polymorphisms within populations, which would require AcLi and acli cyanotypes to be favoured over Acli and acLi cyanotypes (Kooyers and Olsen, 2012).

Our results also do not provide evidence that either directional or epistatic selection would create or maintain cyanogenesis clines across a latitudinal gradient. In fact, our field sampling efforts clearly demonstrate fixation of both *Ac* and *Li* alleles in Louisiana, as expected, while surprisingly, our experimental results suggest that *Ac* alleles should be fixed at both northern and southern study sites. These results are contrary to the LHDH and to previous evidence of exacerbated ecological or allocation costs of HCN production in colder climates as drivers of cyanogenesis cline formation (Daday, 1965; Kooyers et al., 2018). Results of previous studies on deviations from the LHDH do provide some support for our findings, with greater expression of plant defenses in colder climates or plant defenses exhibiting selection patterns based on other environmental factors and/or other plant processes, although these results are often highly variable (Moles et al., 2011; Kooyers et al., 2017; Anstett et al., 2018). It is also possible that rare and stressful extreme climatic events that did not occur during the time period of our experiment select for *ac* and *li* alleles, which could be at least partially responsible for the maintenance of the cyanogenesis polymorphism within most natural *T. repens* populations and cyanogenesis cline formation. Taken together, these findings indicate that understanding the formation of clines in plant defense across complex environmental gradients requires a multifaceted approach to understanding the costs, benefits, and trade-offs associated with plant defenses among populations.

### Limitations

The clearest limitation of our study is the observed lack of effectiveness of the precipitation manipulation in significantly reducing soil moisture availability compared with control subplots at both study sites in 2021. This restricts our ability to detect effects of precipitation manipulation on *T. repens* fitness and to assess precipitation as an agent of selection on cyanogenesis and its metabolic components in *T. repens*. Rainout shelters were modified after the 2021 growing season to reduce precipitation more substantially in the treated subplots and limit the amount of unintentional precipitation reduction in the control subplots. These modifications resulted in significant differences in soil moisture between precipitation reduction subplots and control subplots in Ontario in 2022, but not in Louisiana, likely because of uncharacteristically high rainfall and soil moisture across the growing season at this study site (**Table S1**). However, despite the ineffectiveness of the treatment, in instances where precipitation reduction did have significant effects on *T. repens* fitness in Ontario in 2022, we did not find evidence of the precipitation manipulation imposing differential selection on *T. repens* cyanotypes. This result increases confidence in our evidence against precipitation as a driver of the formation of cyanogenesis clines in *T. repens*, despite mixed effectiveness of the precipitation manipulation.

An additional potential limitation of our study is that individuals were not transplanted into the field until after eight weeks of growth in a common greenhouse environment. This was necessary to provide sufficient time for each seedling’s cyanotype to be confirmed through Feigl-Anger assays before transplanting one individual of each cyanotype into each subplot (Feigl and Anger, 1966). However, the seedling stage is critical to lifetime fitness due to strong viability selection (Agrawal et al., 2013; Barton and Hanley, 2013; Warvell and Shaw, 2019), which could be particularly true for seedling mortality caused by our variables of interest, precipitation and herbivory. The current study experienced very low mortality overall, while a separate experiment transplanting *T. repens* individuals into similar common garden sites after only four weeks experienced substantially higher mortality of seedlings (Albano et al., 2024). Future studies directly investigating the effects of precipitation and herbivory on white clover seedlings across latitudes would be useful to corroborate our results with data across the entire life cycle of *T. repens*.

## Conclusions

We employed a manipulative experiment in natural conditions to investigate precipitation and herbivory as potential agents of natural selection on cyanogenesis, driving the evolution of cline formation at *Ac*/*ac* and *Li*/*li* in *T. repens*. Overall, there was limited evidence for either of these manipulated environmental variables in imposing differential selection on *T. repens* cyanotypes at either study site. Further, in isolated instances that did indicate differential selection, cyanotypes were not affected in a manner consistent with the maintenance of cyanogenesis polymorphisms within populations or clines across latitudinal gradients. Therefore, our results do not support previous hypotheses surrounding precipitation and herbivory as influential factors in the formation of latitudinal cyanogenesis clines. Instead, we found that plants that produce CNglcs, exhibited higher fitness than other cyanotypes in general, regardless of cyanogenesis, and that this effect was stronger at the higher latitude garden, which is again, contrary to expectations based on observed cyanogenesis clines. This indicates that the ability to produce HCN as an antiherbivore defense could still outweigh any potential costs to the plant, even in environments that experience seasonal freezing temperatures that exacerbate costs (Kooyers et al., 2018). Furthermore, the ability to produce CNglcs may provide additional benefits even in the absence of plant-derived β-glycosidases, such as downstream antiherbivore effects through the activity of herbivore-derived β-glycosidases or the ability to recycle N from CNglcs for use in other plant functions. However, direct tests of the significance of these potential benefits are needed in *T. repens* to provide insight into their relative impact on plant fitness. Additionally, future experiments using QTL mapping or genome editing to more definitively isolate the effects of *Ac*/*ac* and *Li*/*li* on plant fitness and environmental response can help elucidate their potentially multifaceted roles in plant function, cline formation, and adaptation to environmental change.

## Supporting information

Albano_et_al_2025_SupplementaryMaterial

## Acknowledgements

We thank Kate Brown, Andre Daugereaux, Peter Duggan, Hayden Fargo, John Jensen, Olivia Toth, and Yimin Yu for assistance with setting up the experiment. We thank Barry Albano, Paula Albano, Samer El-Galmady, Hayden Fargo, Alanah Joyce, Kavya Manikonda, Sophie Mosler, Nial Navaratne, Olivia Toth, Vanessa Trinca, and Yimin Yu for assistance with conducting the field experiment. We thank Paula Albano, Sarah Albano, Keerath Bhachu, Oliver Du, Giuseppe Labricciosa, Lidia Labricciosa, Zain Nasrullah, Nial Navaratne, Narindra Persaud, and Van Vuong for assistance with sample processing. This research was supported by an NSERC Discovery Grant, Canada Research Chair, and NSERC EWR Steacie Fellowship to MTJJ, NSF grants OIA-1920858 and DBI-2244712 to NJK, and an NSERC CGS Doctoral Award to LJA.

## Data Availability Statement

The dataset generated and analyzed during this study is published in Dryad Digital Repository: https://doi.org/10.5061/dryad.4xgxd25n1.

