## Supplementary material for "A field experiment assessing the roles of drought, herbivory, and local climate on cyanogenesis cline formation and local adaptation in *Trifolium repens*": Albano_et_al_2025_SupplementaryMaterial

### Supplementary Methods

*Field surveys*

To test for the presence or absence of latitudinal clines in herbivory and HCN frequency, seeds were harvested and herbivory estimates were taken from 50 natural *T. repens* populations along a 2500 km north-south transect in North America, from Lafayette, Louisiana, USA to Moosonee, Ontario, Canada (~21° span in latitude). Herbivory was estimated visually as percent leaf area consumed for 50 individuals per population (1 trifoliate leaf per individual), following the protocol of Johnson et al. (2016), and then averaged for each population prior to statistical analysis. All leaves sampled were a minimum distance of 3 m from each other to minimize the likelihood of any of the 50 leaves originating from the same clonal mat. The latitude of each sampled population was recorded and mean annual temperature data for each population was extracted from WorldClim at a resolution of 30 seconds (O'Donnell and Ignizio, 2012). Harvested *T. repens* seeds (up to 20 per population) were established in common growth chamber conditions (25°C day, 15°C night, 14 h:10 h light:dark cycle) and hand-pollinated haphazardly within each population to create an F1 generation of seeds. The F1 individuals were transplanted into a common garden field experiment in Mississauga, Ontario, Canada, and after ~1 month of growth in the field, we harvested three trifoliate leaves from each plant for cyanotyping. Estimates of the proportion of cyanogenic phenotypes (AcLi) vs acyanogenic phenotypes (Acli, acLi, and acli) within each population were made from the cyanotypes of 140 plants across 47 populations (3 populations were omitted due to lack of germination success), which were determined using a Feigl-Anger assay (see further details below; Feigl and Anger, 1966).

All statistical analyses were conducted in R Version 4.4.1 (2024) and R Studio Version 2024.09.1 (R Development Core Team, 2024). We constructed three linear models to assess the presence of latitudinal clines in cyanogenesis and herbivory. First, the model: *Mean Annual Temperature* ~ *Latitude* was conducted using the *lm* function fit to a normal distribution. Second, the model: *Cyanogenesis* ~ *Mean Annual Temperature*, was conducted using the *glm* function fit to a binomial distribution with a logit link function. Third, the model: *Herbivory* ~ *Mean Annual Temperature*, was conducted using the *lm* function fit to a normal distribution. For models constructed using the *lm* function, $r^2$ and $p$ values were calculated using the *summary* function. For models constructed using the *glm* function, $\chi^2$ and $p$ values were calculated using the *Anova* function in the *car* package (Version 3.1-2; Fox and Weisberg, 2019).

*Experimental plant generation*

Plants of each generation, from the original parents to F3, were grown, maintained, and crossed in the same growth chamber conditions (25°C day, 15°C night, 14 h:10 h light:dark cycle). Our crossing procedure was performed to randomize the dominant and recessive alleles from each locus (*Ac*/*ac* and *Li*/*li*) onto a common genetic background. Past studies have been limited by not isolating the *Ac*/*ac* and *Li*/*li* alleles in this manner (but see Kooyers et al., 2014; Kooyers et al., 2018), making it difficult to discern selection on these specific loci versus linked genes. The F1 generation was created by crossing an individual of an *AcAcLiLi* genotype originating from Lafayette, Louisiana, with an individual of an *acaclili* genotype originating from King Township, Ontario. Phenotyping 46 F1 progeny from this cross showed that all plants were cyanogenic and therefore double heterozygotes (*AcacLili*). Once flowering occurred for F1 individuals, each available inflorescence was pollinated with every other available inflorescence to obtain an F2 generation of seeds, with cyanotypes expected to be present in a 9:3:3:1 ratio (*Ac_Li_* : *Ac_li_* : *ac_Li_* : *ac_li_*). The F2 plants were then separated by cyanotype and haphazardly outcrossed via hand-pollination within each cyanotype to obtain an F3 generation of seeds for use in the experiment.

The cyanotype of each F3 individual selected for the experiment was confirmed using three sets of Feigl-Anger assays (Feigl and Anger, 1966) on macerated *T. repens* leaf tissue as follows: (1) to test for the presence/absence of HCN, leaf tissue was macerated in 80 μL of deionized water, with positive tests indicating AcLi plants; (2) to test for the presence/absence of CNglcs, leaf tissue was macerated in 80 μL of 0.2 EU/mL β-glucosidase (i.e., linamarase; Sigma Aldrich, Oakville, Ontario, Canada) and 20 μL of deionized water, with positive tests indicating Acli plants; (3) to test for the presence/absence of linamarase, leaf tissue was macerated in 15 μL of 10 mM linamarin (Sigma Aldrich, Oakville, Ontario, Canada) and 65 μL of deionized water, with positive tests indicating acLi plants. All plants that had a negative test in all three assays were determined to be lacking both CNglcs and linamarase (acli).

### Supplementary Tables and Figures

**Table S1** Historical (1971-2000) and current (during the years of the study; 2021 and 2022) climatic characteristics and elevation at each study site. Data for mean annual temperature, mean temperature of the coldest month, mean temperature of the warmest month, annual precipitation, Hargreaves climatic moisture deficit, and Hargreaves reference evaporation (potential evapotranspiration) were extracted from ClimateNA (Wang et al., 2016). Aridity index was calculated as the quotient of annual precipitation and potential evapotranspiration for each time period.

| | Ontario | | | Louisiana | | |
|---|---|---|---|---|---|---|
| Variable | Historical | 2021 | 2022 | Historical | 2021 | 2022 |
| Mean Annual Temperature (°C) | 7.7 | 9.8 | 8.5 | 19.8 | 20.1 | 20.0 |
| Mean Temperature (Coldest Month; °C) | −6.2 | −6.3 | −8.9 | 10.4 | 9.3 | 9.5 |
| Mean Temperature (Warmest Month; °C) | 20.7 | 22.9 | 21.8 | 27.8 | 27.9 | 28.4 |
| Annual Precipitation (mm) | 740 | 881 | 751 | 1508 | 2085 | 1463 |
| Climatic Moisture Deficit (mm) | 229 | 216 | 268 | 158 | 55 | 291 |
| Potential Evapotranspiration (mm) | 704 | 779 | 789 | 1380 | 1296 | 1387 |
| Aridity Index | 1.051 | 1.131 | 0.952 | 1.093 | 1.609 | 1.055 |
| Elevation (m) | 283 | | | 13 | | |

**Table S2** Soil moisture measurements (in percent volume), analyzed separately for the 2021 and 2022 growing seasons at the Ontario and Louisiana study sites. Mixed effects models were analyzed using the function *lmer* in the *lme4* package (Version 1.1-27.1; Bates et al., 2014) and the following model structure: % Volume ~ Precipitation + Sampling Date + Precipitation:Sampling Date + (1|Whole Plot ID/Subplot ID). Precipitation (control versus reduced) and sampling date were included as fixed effects, while subplot ID nested within whole plot ID was included as a random effect to account for non-independence among the replicate measurements taken in each subplot and non-independence of the precipitation manipulation among whole plots. Significant effects ($P$ < 0.05) are listed in bold.

| **Ontario Study Site** | 2021 | | | 2022 | | |
|---|---|---|---|---|---|---|
| Predictor | $\chi^2$ | df | $P$ | $\chi^2$ | df | $P$ |
| Precipitation | 0.447 | 1 | 0.504 | **373.373** | **1** | **< 0.001** |
| Sampling Date | **355.589** | **1** | **< 0.001** | **645.888** | **5** | **< 0.001** |
| Precipitation × Sampling Date | 0.023 | 1 | 0.878 | **80.028** | **5** | **< 0.001** |

| **Louisiana Study Site** | 2021 | | | 2022 | | |
|---|---|---|---|---|---|---|
| Predictor | $\chi^2$ | df | $P$ | $\chi^2$ | df | $P$ |
| Precipitation | 0.642 | 1 | 0.423 | 0.040 | 1 | 0.841 |
| Sampling Date | **1153.63** | **19** | **< 0.001** | **402.142** | **5** | **< 0.001** |
| Precipitation × Sampling Date | 15.948 | 19 | 0.661 | **21.822** | **5** | **< 0.001** |

**Table S3** Summary of transformations used for each quantitative response variable. Transformations were performed to meet assumptions of normality and homogeneity of variance. All models described in the table are fit to a normal distribution. All models in the study that were fit to the binomial distribution (survival and whether plants produced flowers and/or seeds) did not require further transformation so are not included in the table.

| Response | # of Flower Heads | Seed Set Mass | Maximum Plant Area | Growth Rate | Herbivory |
|---|---|---|---|---|---|
| Ontario 2021 | Square root | Square root | Square root | None | Log |
| Ontario 2022 | Square root | Square root | Square root | Square root | Log |
| Louisiana 2021 | Log | Log | Square root | Square root | Log |
| Louisiana 2022 | Log | Log | Square root | Log | Log |

**Table S4** Results of mixed effects models assessing how cyanotype, manipulation of precipitation, and manipulation of herbivores affected *T. repens* growing in Ontario in the 2021 growing season. Response variables include (winter) survival, whether plants produced flowers, whether plants produced seeds, maximum plant area, growth rate, and herbivory. Models for survival did not converge so this variable was replaced by winter survival in this table. Signs of > and < are with respect to the first listed side of the custom contrast (cyanogenic, *Ac*, or *Li*). Significant effects ($P$ < 0.05) are listed in bold.

| | Winter Survival | | | # of Plants Producing Flowers | | | # of Flower Heads | | |
|---|---|---|---|---|---|---|---|---|---|
| Predictor | $\chi^2$ | df | $P$ | $\chi^2$ | df | $P$ | $\chi^2$ | df | $P$ |
| Cyanotype (C) | 0.901 | 3 | 0.825 | 5.540 | 3 | 0.136 | **10.476** | **3** | **0.015** |
| Precipitation (P) | 0.137 | 1 | 0.711 | 0.002 | 1 | 0.961 | 0.042 | 1 | 0.838 |
| Herbivores (H) | 3.007 | 1 | 0.083 | 1.533 | 1 | 0.216 | 0.075 | 1 | 0.784 |
| C × P | 2.852 | 3 | 0.415 | 5.870 | 3 | 0.118 | 4.427 | 3 | 0.219 |
| C × H | 1.649 | 3 | 0.648 | 1.541 | 3 | 0.673 | 1.602 | 3 | 0.659 |
| P × H | 0.523 | 1 | 0.470 | 0.418 | 1 | 0.518 | 0.226 | 1 | 0.635 |
| Contrast | Sign | | $P$ | Sign | | $P$ | Sign | | $P$ |
| *Cyanotype* | | | | | | | | | |
| Cyanogenic vs. Acyanogenic | — | | — | — | | — | > | | **0.010** |
| *Ac* vs. *ac* | — | | — | — | | — | > | | **0.014** |
| *Li* vs. *li* | — | | — | — | | — | = | | 0.955 |

| | Maximum Plant Area | | | Growth Rate | | | Herbivory | | |
|---|---|---|---|---|---|---|---|---|---|
| Predictor | $\chi^2$ | df | $P$ | $\chi^2$ | df | $P$ | $\chi^2$ | df | $P$ |
| Cyanotype (C) | 3.319 | 3 | 0.345 | 2.000 | 3 | 0.572 | **55.384** | **3** | **< 0.001** |
| Precipitation (P) | 0.985 | 1 | 0.321 | 0.005 | 1 | 0.941 | 0.039 | 1 | 0.843 |
| Herbivores (H) | 3.138 | 1 | 0.076 | **18.350** | **1** | **< 0.001** | **61.480** | **1** | **< 0.001** |
| C × P | 1.430 | 3 | 0.699 | 0.224 | 3 | 0.974 | 2.978 | 3 | 0.395 |
| C × H | 3.025 | 3 | 0.388 | 2.071 | 3 | 0.558 | 4.905 | 3 | 0.179 |
| P × H | 1.932 | 1 | 0.165 | 0.089 | 1 | 0.765 | 0.658 | 1 | 0.417 |
| Contrast | Sign | | $P$ | Sign | | $P$ | Sign | | $P$ |
| *Cyanotype* | | | | | | | | | |
| Cyanogenic vs. Acyanogenic | — | | — | — | | — | < | | **< 0.001** |
| *Ac* vs. *ac* | — | | — | — | | — | < | | **< 0.001** |
| *Li* vs. *li* | — | | — | — | | — | = | | 0.977 |

**Table S5** Results of mixed effects models assessing how cyanotype, manipulation of precipitation, and manipulation of herbivores affected *T. repens* growing in Louisiana in the 2021 growing season. Response variables include survival, whether plants produced flowers, whether plants produced seeds, maximum plant area, growth rate, and herbivory. Signs of > and < are with respect to the first listed side of the custom contrast (cyanogenic, *Ac*, or *Li*). Significant effects ($P < 0.05$) are listed in bold.

| | Survival | | | # of Plants Producing Flowers | | | # of Flower Heads | | |
|---|---|---|---|---|---|---|---|---|---|
| Predictor | $\chi^2$ | df | *P* | $\chi^2$ | df | *P* | $\chi^2$ | df | *P* |
| Cyanotype (C) | 2.028 | 3 | 0.567 | **10.874** | **3** | **0.012** | 3.600 | 3 | 0.308 |
| Precipitation (P) | 1.752 | 1 | 0.186 | 1.919 | 1 | 0.166 | 1.273 | 1 | 0.259 |
| Herbivores (H) | 0.000 | 1 | 0.984 | **16.043** | **1** | **< 0.001** | **44.758** | **1** | **< 0.001** |
| C × P | 1.171 | 3 | 0.760 | 0.613 | 3 | 0.894 | 0.982 | 3 | 0.806 |
| C × H | 2.552 | 3 | 0.466 | 1.817 | 3 | 0.611 | 3.651 | 3 | 0.302 |
| P × H | 0.000 | 1 | 0.997 | 0.001 | 1 | 0.978 | 1.947 | 1 | 0.163 |
| Contrast | Sign | | *P* | Sign | | *P* | Sign | | *P* |
| *Cyanotype* | | | | | | | | | |
| Cyanogenic vs. Acyanogenic | — | | — | > | | **0.003** | — | | — |
| *Ac* vs. *ac* | — | | — | > | | **0.037** | — | | — |
| *Li* vs. *li* | — | | — | > | | **0.002** | — | | — |

| | Maximum Plant Area | | | Growth Rate | | | Herbivory | | |
|---|---|---|---|---|---|---|---|---|---|
| Predictor | $\chi^2$ | df | *P* | $\chi^2$ | df | *P* | $\chi^2$ | df | *P* |
| Cyanotype (C) | **13.424** | **3** | **0.004** | 7.546 | 3 | 0.056 | 1.848 | 3 | 0.605 |
| Precipitation (P) | 0.149 | 1 | 0.700 | 0.000 | 1 | 0.988 | 1.079 | 1 | 0.299 |
| Herbivores (H) | **12.782** | **1** | **< 0.001** | **7.886** | **1** | **0.005** | **186.90** | **1** | **< 0.001** |
| C × P | 1.518 | 3 | 0.678 | 3.610 | 3 | 0.307 | 1.275 | 3 | 0.735 |
| C × H | 5.642 | 3 | 0.130 | 3.649 | 3 | 0.302 | 1.271 | 3 | 0.736 |
| P × H | 0.346 | 1 | 0.557 | 1.017 | 1 | 0.313 | 2.432 | 1 | 0.119 |
| Contrast | Sign | | *P* | Sign | | *P* | Sign | | *P* |
| *Cyanotype* | | | | | | | | | |
| Cyanogenic vs. Acyanogenic | = | | 0.089 | — | | — | — | | — |
| *Ac* vs. *ac* | = | | 0.770 | — | | — | — | | — |
| *Li* vs. *li* | = | | 0.949 | — | | — | — | | — |

**Table S6** Results of mixed effects models assessing how cyanotype, manipulation of precipitation, and manipulation of herbivores affected *T. repens* growing in Ontario in the 2022 growing season. Response variables include survival, whether plants produced flowers, whether plants produced seeds, maximum plant area, growth rate, and herbivory. Signs of > and < are with respect to the first listed side of the custom contrast (cyanogenic, *Ac*, or *Li*). Significant effects ($P < 0.05$) are listed in bold.

| | Survival | | | # of Plants Producing Flowers | | | # of Flower Heads | | |
|---|---|---|---|---|---|---|---|---|---|
| Predictor | $\chi^2$ | df | *P* | $\chi^2$ | df | *P* | $\chi^2$ | df | *P* |
| Cyanotype (C) | **21.666** | **3** | **< 0.001** | **18.015** | **3** | **< 0.001** | **13.122** | **3** | **0.004** |
| Precipitation (P) | 0.001 | 1 | 0.979 | 1.505 | 1 | 0.220 | 0.844 | 1 | 0.358 |
| Herbivores (H) | **7.495** | **1** | **0.006** | **4.679** | **1** | **0.031** | **7.145** | **1** | **0.008** |
| C × P | 3.325 | 3 | 0.344 | 7.460 | 3 | 0.059 | 2.321 | 3 | 0.509 |
| C × H | 7.378 | 3 | 0.061 | 2.402 | 3 | 0.493 | 0.664 | 3 | 0.882 |
| P × H | **5.116** | **1** | **0.024** | 3.035 | 1 | 0.081 | 3.707 | 1 | 0.054 |
| Contrast | Sign | | *P* | Sign | | *P* | Sign | | *P* |
| *Cyanotype* | | | | | | | | | |
| Cyanogenic vs. Acyanogenic | > | | **0.006** | = | | 0.104 | > | | **0.004** |
| *Ac* vs. *ac* | > | | **< 0.001** | > | | **< 0.001** | > | | **0.002** |
| *Li* vs. *li* | = | | 0.618 | = | | 0.085 | = | | 0.812 |

| | Maximum Plant Area | | | Growth Rate | | | Herbivory | | |
|---|---|---|---|---|---|---|---|---|---|
| Predictor | $\chi^2$ | df | *P* | $\chi^2$ | df | *P* | $\chi^2$ | df | *P* |
| Cyanotype (C) | **25.443** | **3** | **< 0.001** | **10.074** | **3** | **0.018** | **25.248** | **3** | **< 0.001** |
| Precipitation (P) | 3.806 | 1 | 0.051 | 0.180 | 1 | 0.671 | 0.043 | 1 | 0.836 |
| Herbivores (H) | **14.908** | **1** | **< 0.001** | 0.272 | 1 | 0.602 | **29.722** | **1** | **< 0.001** |
| C × P | 3.685 | 3 | 0.298 | 0.802 | 3 | 0.849 | 6.014 | 3 | 0.111 |
| C × H | 2.827 | 3 | 0.419 | 0.343 | 3 | 0.952 | 2.263 | 3 | 0.520 |
| P × H | 0.148 | 1 | 0.700 | 1.133 | 1 | 0.287 | 0.636 | 1 | 0.425 |
| Contrast | Sign | | *P* | Sign | | *P* | Sign | | *P* |
| *Cyanotype* | | | | | | | | | |
| Cyanogenic vs. Acyanogenic | > | | **0.002** | < | | **0.002** | < | | **< 0.001** |
| *Ac* vs. *ac* | > | | **< 0.001** | > | | **0.009** | < | | **< 0.001** |
| *Li* vs. *li* | = | | 0.643 | = | | 0.643 | = | | 0.982 |

**Table S7** Results of mixed effects models assessing how cyanotype, manipulation of precipitation, and manipulation of herbivores affected *T. repens* growing in Louisiana in the 2022 growing season. Response variables include survival, whether plants produced flowers, whether plants produced seeds, maximum plant area, growth rate, and herbivory. Signs of > and < are with respect to the first listed side of the custom contrast (cyanogenic, *Ac*, or *Li*). Significant effects ($P < 0.05$) are listed in bold.

| | Survival | | | # of Plants Producing Flowers | | | # of Flower Heads | | |
|---|---|---|---|---|---|---|---|---|---|
| Predictor | $\chi^2$ | df | *P* | $\chi^2$ | df | *P* | $\chi^2$ | df | *P* |
| Cyanotype (C) | 2.340 | 3 | 0.505 | 0.982 | 3 | 0.806 | **13.100** | **3** | **0.004** |
| Precipitation (P) | 3.715 | 1 | 0.054 | 1.586 | 1 | 0.208 | 0.860 | 1 | 0.354 |
| Herbivores (H) | 2.556 | 1 | 0.110 | 0.013 | 1 | 0.909 | 0.048 | 1 | 0.826 |
| C × P | 6.844 | 3 | 0.077 | 3.988 | 3 | 0.263 | 3.451 | 3 | 0.327 |
| C × H | 4.524 | 3 | 0.210 | 0.076 | 3 | 0.995 | 1.195 | 3 | 0.754 |
| P × H | 1.594 | 1 | 0.207 | 0.187 | 1 | 0.666 | 1.318 | 1 | 0.251 |
| Contrast | Sign | | *P* | Sign | | *P* | Sign | | *P* |
| *Cyanotype* | | | | | | | | | |
| Cyanogenic vs. Acyanogenic | — | | — | — | | — | = | | 0.537 |
| *Ac* vs. *ac* | — | | — | — | | — | > | | **0.002** |
| *Li* vs. *li* | — | | — | — | | — | = | | 0.529 |

| | Maximum Plant Area | | | Growth Rate | | | Herbivory | | |
|---|---|---|---|---|---|---|---|---|---|
| Predictor | $\chi^2$ | df | *P* | $\chi^2$ | df | *P* | $\chi^2$ | df | *P* |
| Cyanotype (C) | 5.626 | 3 | 0.131 | 4.365 | 3 | 0.225 | 0.293 | 3 | 0.961 |
| Precipitation (P) | 0.125 | 1 | 0.724 | **4.287** | **1** | **0.038** | 0.019 | 1 | 0.890 |
| Herbivores (H) | **6.338** | **1** | **0.012** | 0.006 | 1 | 0.940 | 2.172 | 1 | 0.141 |
| C × P | 3.209 | 3 | 0.361 | 6.052 | 3 | 0.109 | 5.429 | 3 | 0.143 |
| C × H | 0.714 | 3 | 0.870 | 3.821 | 3 | 0.282 | **8.473** | **3** | **0.037** |
| P × H | 2.171 | 1 | 0.141 | 0.802 | 1 | 0.371 | 0.131 | 1 | 0.718 |
| Contrast | Sign | | *P* | Sign | | *P* | Sign | | *P* |
| *Cyanotype* | | | | | | | | | |
| Cyanogenic vs. Acyanogenic | — | | — | — | | — | — | | — |
| *Ac* vs. *ac* | — | | — | — | | — | — | | — |
| *Li* vs. *li* | — | | — | — | | — | — | | — |

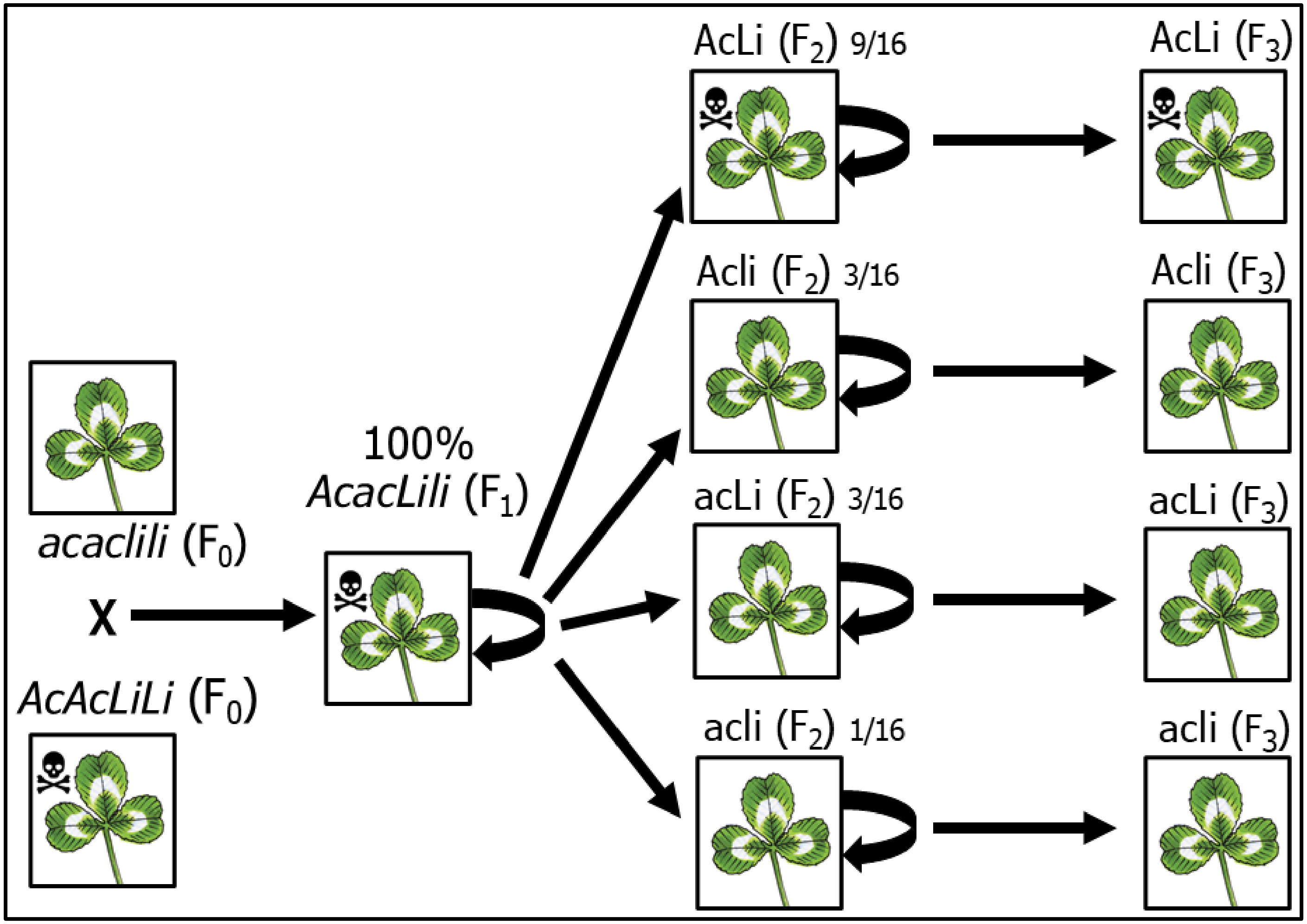

**Figure S1** F3 crossing design used to generate all *T. repens* individuals for this experiment. One *AcAcLiLi* F0 parent from Louisiana was crossed with one *acaclili* F0 parent from Ontario to create an F1 generation of 100% *AcacLili* offspring. All flowering F1 offspring were crossed haphazardly with each other to produce and F2 generation containing all four *T. repens* cyanotypes in normal segregating ratios. All flowering F2 offspring were crossed haphazardly within cyanotype to produce an F3 generation of all four *T. repens* cyanotypes while randomizing the rest of the genetic background. The F3 generation seeds were grown in a common environment while cyanotypes were confirmed using Feigl-Anger assays (Feigl and Anger, 1966) before being transplanted into both study sites.

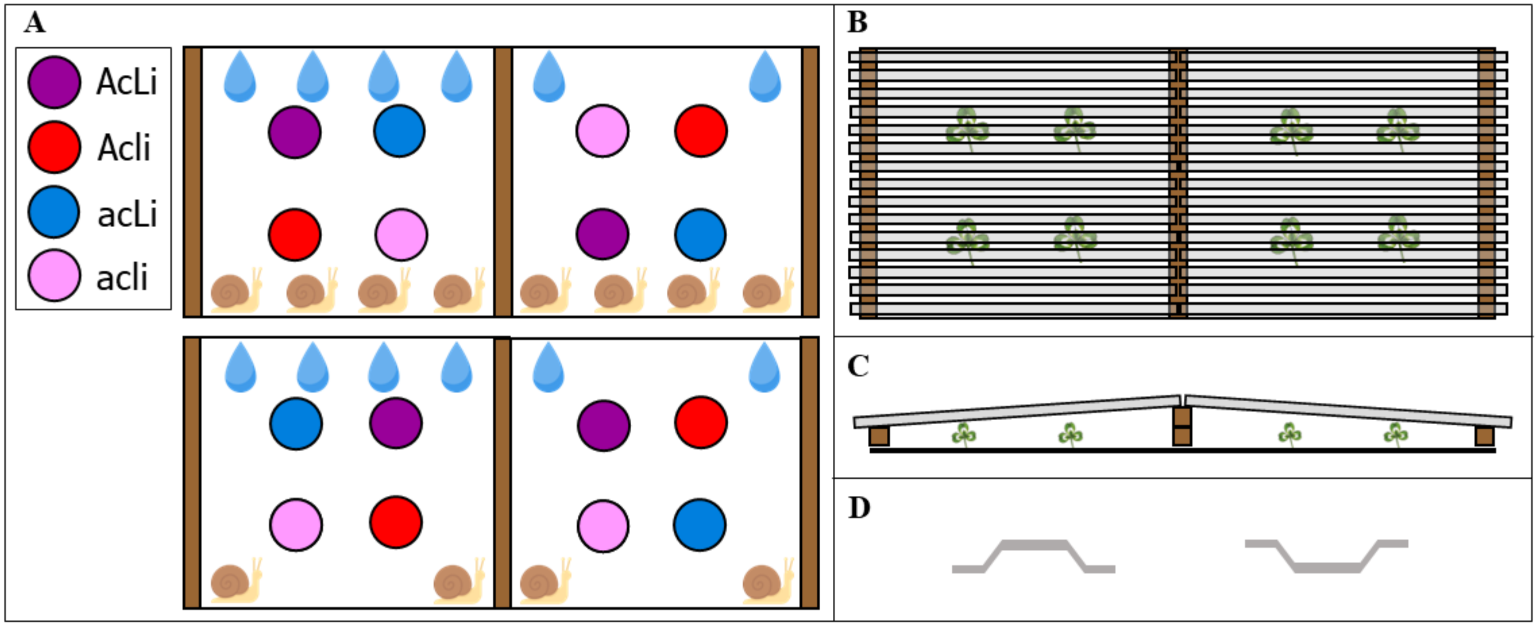

**Figure S2** Diagrams of the split-plot experimental design and the design of the rainout shelters (in Ontario). (A) Top-down view of two example whole plots, each divided into two subplots. Subplot coloration represents the four different treatment combinations in the study. Of the 48 whole plots, 24 were randomly selected for herbivore reduction (bottom) while the other 24 remained as controls (top). Within each whole plot, one subplot was assigned for precipitation reduction while the other remained as a control. In Louisiana, the combination of precipitation reduction and herbivore reduction treatments (four treatment combinations between the two 2-level manipulations) was applied at the subplot level. Within each subplot, different colours represent the four randomly-arranged individual plants of different *T. repens* cyanotypes. All individuals were spaced 0.5 m from each other and from the edge of the shelter to minimize competition and the likelihood of wind-affected precipitation impacting the treatment. (B) A top-down view of an example whole plot based on the 2021 experimental design, demonstrating the use of rainout shelters to apply the precipitation reduction treatment. Horizontal grey bars represent transparent, corrugated polycarbonate strips attached to each shelter frame. Leaves represent *T. repens* individuals beneath each shelter. (C) A side view of an example whole plot and rainout shelters, demonstrating the use of two stacked 4” × 4” pieces of lumber between the subplots and one 4” × 4” at each end, which provides an angle to the shelters to allow for run-off of intercepted precipitation. (D) End view of transparent, corrugated polycarbonate strips, demonstrating the use of strip orientation to apply the precipitation reduction treatment. For the shelter in each whole plot assigned for precipitation reduction, all strips were oriented with the opening facing upwards (left) to intercept incoming precipitation. For the other shelter in each whole plot, all strips were oriented with the opening facing downwards (right) to allow precipitation to reach the *T. repens* plants.

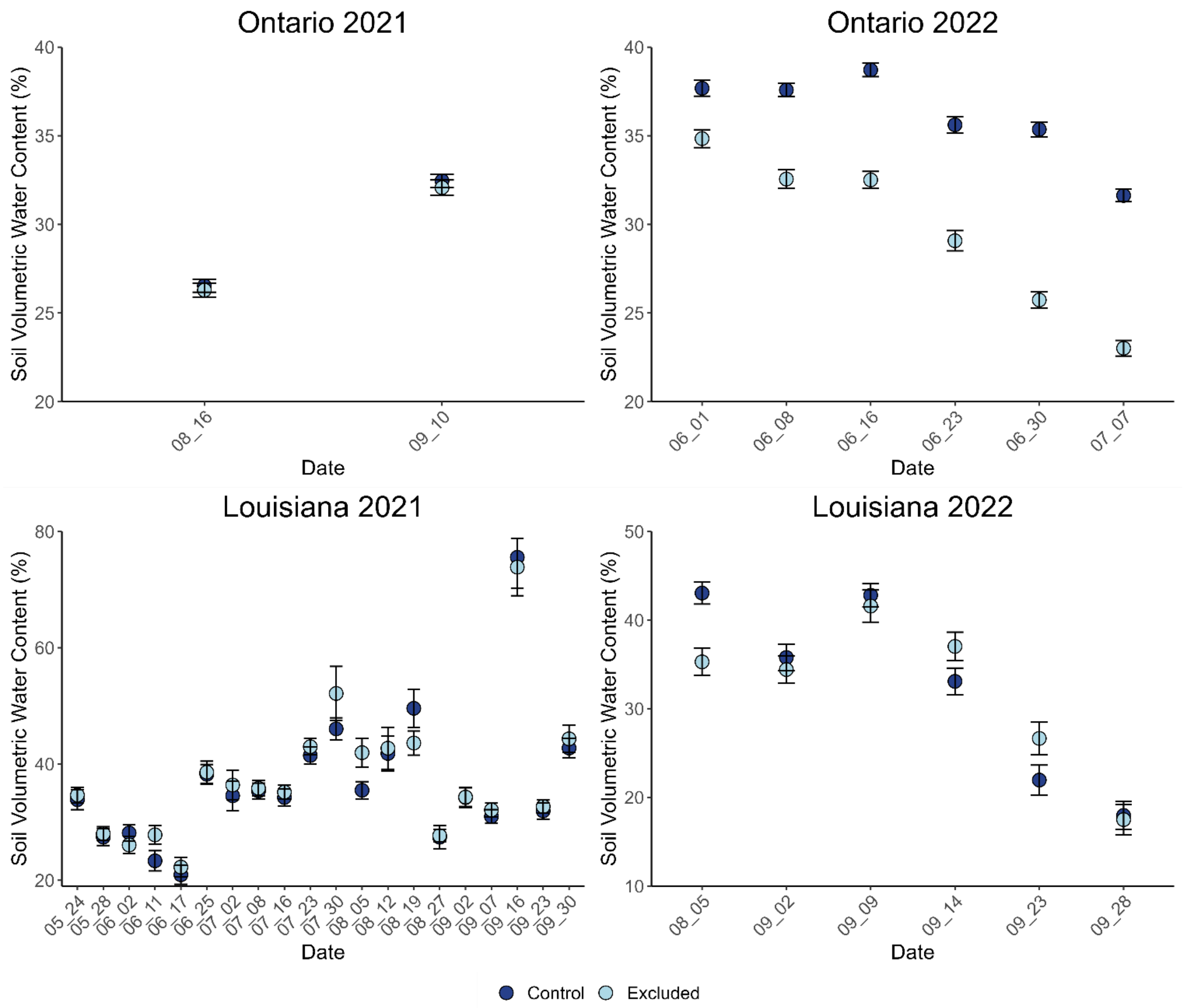

**Figure S3** Soil moisture at each study site in each year of the experiment. We show soil volumetric water content at various dates throughout the 2021 and 2022 growing seasons at both the Ontario and Louisiana study sites. Error bars are ± one standard error for each mean value.
